# *Salmonella* Typhimurium effector SseI regulates host peroxisomal dynamics to acquire lysosomal cholesterol for better intracellular growth

**DOI:** 10.1101/2023.02.27.530266

**Authors:** Desh Raj, Abhilash Vijay Nair, Jyotsna Sharma, Shakti Prakash, Aman Kaushik, Swarnali Basu, Shikha Sahu, Shriya Singh, Vivek Bhosale, Tulika Chandra, Uday C Ghoshal, Arunava Dasgupta, Mohammad Imran Siddiqi, Shashi Kumar Gupta, Dipshikha Chakravortty, Veena Ammanathan, Amit Lahiri

## Abstract

Intracellular *Salmonella* resides and multiplies in cholesterol-rich specialized compartment called *Salmonella*-containing vacuoles (SCVs) and avoids fusion with acidic lysosomes. Given, lysosomes are primary organelle that redistributes LDL derived cholesterol to other organelles; we questioned how lysosomal cholesterol can be transported to SCV. We demonstrate here that peroxisomes are recruited to SCVs in human primary macrophages, epithelial cells and functions as pro-bacterial organelles. Further, this interaction is assisted by SseI, a *Salmonella* effector protein containing mammalian peroxisome targeting sequence. SseI localizes to peroxisome, interacts and activates a host Ras GTPase, ARF-1 on the peroxisome membrane. Activation of ARF-1 leads to recruitment of phosphatidylinsolitol-5- phosphate-4 kinase to generate phosphatidylinsolitol-4-5-bisphosphate on peroxisomes. Accordingly, the *ΔsseI* strain showed reduced virulence in cell lines and during mice infection. Taken together, our work identified a fascinating mechanism by which a pathogen targets host organelles via its secretory effectors and exploits host metabolic intermediates for its intracellular proliferation.

## Introduction

Enteric pathogen *Salmonella enterica* serotype Typhimurium (STM), the causative agent of gastroenteritis in humans, can invade and survive within macrophages and epithelial cells and replicate in distinct membrane-bound vesicles known as *Salmonella*-containing vacuoles (SCVs)^1^. SCVs develop tubular extensions known as *Salmonella* induced filaments (SIFs) that are important for the growth and survival of *Salmonella* in the host cells^2^. Formation and maintenance of SCVs require expression of bacterial effector proteins encoded by type three secretion systems (TTSS) namely *Salmonella* pathogenicity islands 1&2 (SPI-1/2)^3, 4^. SPI1 TTSS is required for entry of the bacteria in the epithelial cells while SPI2 TTSS leads to the secretion of more than 30 proteins across the SCV membrane in both epithelial cells and macrophages^5, 6^. SPI2 TTSS also promotes intracellular replication of STM by altering antibacterial peptide, ROS and NO mediated killing, decreasing lysosomal fusion to SCV, and acquire metabolites required for bacterial growth^7–9^. One of the SPI2 TTSS effector proteins, SseI is a 37 kDa protein and few studies have implicated its role in directing migration of STM infected dendritic cells10, 11, 12.

Cholesterol is shown to accumulate excessively (about 30% of total cellular cholesterol) around the vacuoles/SIF and is essential for SCV/SIF rigidity^13^. SPI2 effector proteins, SseJ and SseL were previously found to the host cholesterol transport protein, oxysterol binding protein 1 (Osbp1) to the cytosolic surface of SCVS. This facilitates non-vesicular transport of cholesterol to SCVS^14^. Further, LDL-r receptor knockout animal are resistant to *Salmonella* infection and human SNP-vac14 in the cholesterol transporter leads to reduced bacterial growth^15, 16^. In addition, *Salmonella* cannot synthesize cholesterol and the sterol that accumulates in SCV is not of biosynthetic origin highlighting need of studies to understand cholesterol transport to SCV for novel host targeted therapy^17–20^. The mechanism by which *Salmonella* acquires cellular cholesterol during active replication inside the SCV is not yet clear.

Recent studies indicate the role of peroxisomes in immune cell activation, inflammation, and cholesterol transport^21–24^. Majority of peroxisomal matrix proteins contain a consensus signal-sequence at the c-terminal (s/a-k/r-l/m) which is known as peroxisomal targeting sequence1 (pts1)^25, 26^. Mapping contacts reveal that peroxisomes are predominantly in contact with ER, mitochondria, lipid droplets and lysosomes ^27, 28^. Extracellular LDL gets transported to lysosomes after binding to its cognate receptor LDL-R and later this LDL derived cholesterol is shuttled to peroxisomes. Peroxisomes redistribute the same to ER by membrane contacts of the lipid PIP2 and tethering protein syt7 between two organelles^24, 29^. The amount of PIP2 level on the organelle can decide organelle movement and the strength of these intra-organellar contact^30^. Interestingly, multiple human proteins including host small GTPase ARF1 regulate PIP2 levels on the peroxisomal membranes by recruiting PIP5 kinases and can directly dictate the cholesterol shuttling^31–33^.

Given SCV do not fuses with lysosomes^34^, we set out to identify how LDL derived cholesterol is conveyed from lysosome to the growing SCV. We have shown in this current work that SCV and the growing SIF associates with peroxisome to assist in *Salmonella* growth in human primary macrophages and in the epithelial cells. We have identified a host pts1 signal in the SPI2 effector protein, SseI and show how SseI can regulate the peroxisomal PIP2 to enhance lysosome-peroxisome and peroxisome-SCV contacts and provide the pathogen a survival advantage in the infected host. As per our knowledge, this is the first report of a bacterial protein having functional human pts1 signal and a remarkable mechanism by which pathogenic bacteria reroutes host lysosomal cholesterol using peroxisome as a bridge.

## Results

Live ***Salmonella*** Typhimurium infection temporally modulates peroxisome dynamics in primary human macrophages and epithelial cells. Since, little is known about the role of peroxisomes during intracellular bacterial infection, we therefore aimed to monitor dynamics of peroxisomes (peroxisomal localization, number and protein expression) during the progression of *Salmonella* Typhimurium(14028S) infection. Interestingly, temporal interaction of peroxisomes (using peroxisome membrane protein pex-14) with SCVs and SIFs marked by LAMP1 (a marker of SCV) and whole bacteria was observed in primary human intestinal epithelial cells isolated from human colon biopsy samples (Fig.1A). We next used human blood derived primary macrophages and macrophage cell line THP1 and similar peroxisome temporalities were observed (Fig.1B). To better understand the significance of this association, human primary macrophages were infected for 6 hours with either heat killed *Salmonella*, latex bead *or Mycobacterium* smegmatis, *E. coli*. The interaction between peroxisomes was lost in all these conditions in comparison with wild type *Salmonella* suggesting that only live virulent *Salmonella* induces this interaction (Supplementary figure 1A-D). We utilized Human Epithelial cell line HeLa in the following experiments and observed starting from 3 hour of infection WT *Salmonella* showed significant colocalization with peroxisomes and percent colocalization of ‘LAMP-1 positive SCV’ with Pex14 increased over time till 12 hours (Fig1C-D). Viruses like HCMV, HSV-1, SARS-CoV-2 lead to increase in peroxisome numbers late in infection with a concomitant increase in the peroxisomal protein level^35, 36^. Volcano plot from our proteomic analysis of *Salmonella* infected HeLa cells showed no change in the identified 40 peroxisome associated proteins (out of total 3412 proteins) like for fission (PEX11b), de novo biogenesis (PEX3, PEX19), membrane (PEX14, ABCD1), and matrix (catalase) (Fig. 2A). In accordance with this, the peroxisome numbers (Fig.2B) or enzyme activities (Fig.2C) were not altered after *Salmonella* infection in spite of the observed colocalization. We validated the proteomics results by doing RNA expression studies and found that peroxisome biogenesis (PEX19), fission (PEX11B) and membrane proteins like PEX14 expression were not altered after 6 hr of *Salmonella* infection (Supplementary Fig.2A). Next, protein expression of PEX14 was analyzed and *Salmonella* infection did not increase PEX14 translation after infection in the HeLa cells (Supplementary Fig.2B). Taken together, these results indicate temporal increase in the interaction of peroxisome with at early time point after *S.* Typhimurium infection without any change in peroxisome dynamics and metabolism.

**Figure 1.**
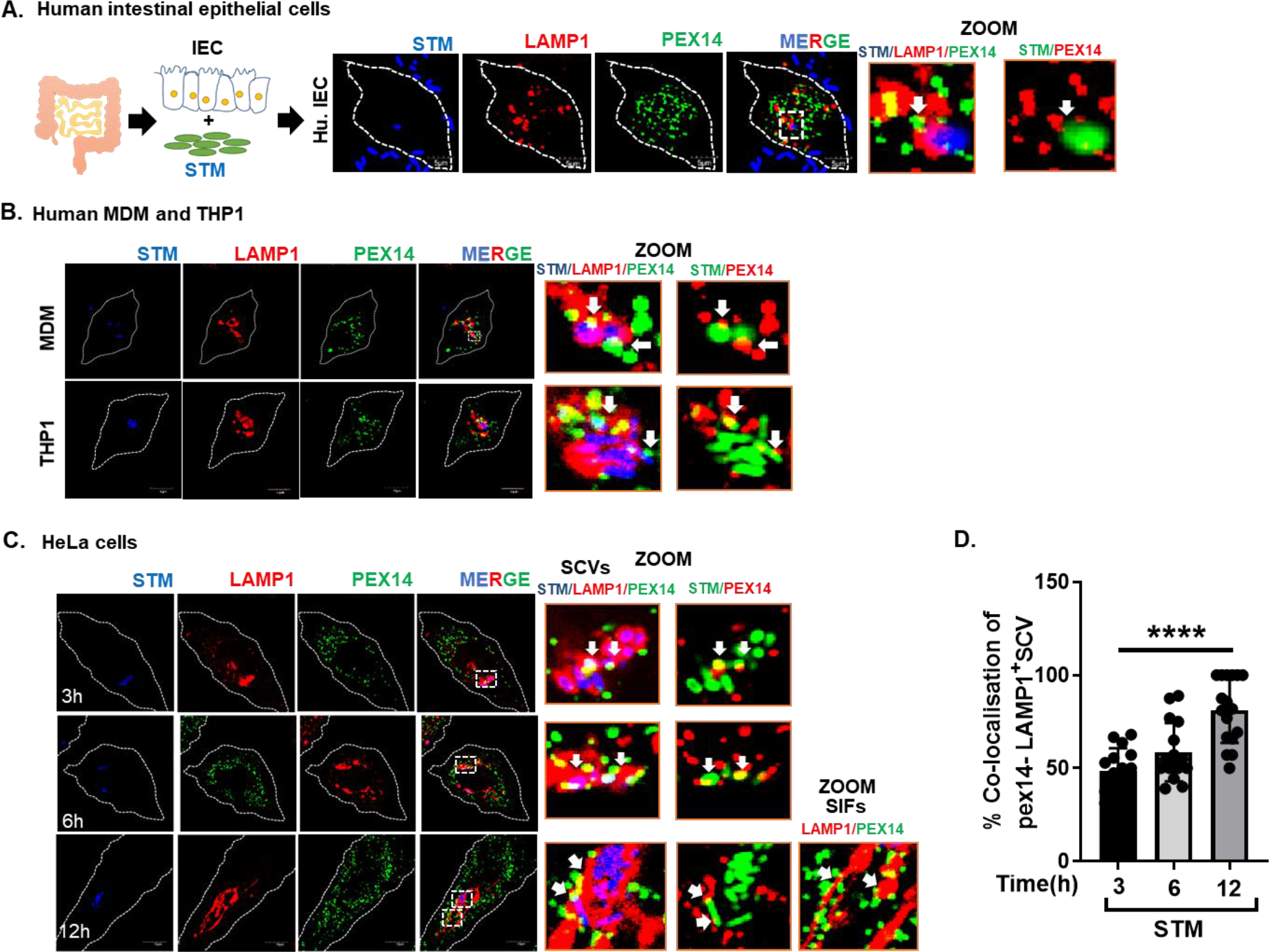
***Salmonella*** Typhimurium (STM) infection induces peroxisome interaction with SCV temporally. A. Representative microscopy images of primary human intestinal epithelial cells isolated from colonic biopsy sample were infected with STM (MOI=50) for 3 hours to check colocalization with peroxisome. Cells were fixed; stained for PEX14 (green), LAMP1 (red), and STM (blue); and Zoom part of the image are shown (white arrow) colocalization pex14(green) with SCV(LAMP1) and pex14(red) with STM (green); scale bar 5 um. B. Human monocyte derived macrophages (MDM) and THP1 cells were infected with STM (MOI=50) for 6 hours to check colocalization with peroxisome. Cells were fixed; stained for PEX14(green), LAMP1(red), and STM (blue); and Zoom part of image were showing (white arrow) colocalization pex14(green) with SCV(LAMP1) and pex14(red) with STM (green); scale bar 10 um. C. HeLa cells were infected and stained with Pex14 (green) and LAMP1 (red) after infection with STM (blue) at indicated time points (3hr, 6hr, 12hr). White arrow denoted co-localization (yellow) between Pex14-LAMP1 and Pex14-STM(GFP). Similar results were found in three independent experiments; scale bar 10 um. D. Co-localization percentage of Pex14-LAMP1^+^ SCV was quantitated (n=20 cells). Individual data points represent mean±SD. Result is representative of 3 independent experiments. ‘MOI’ denotes multiplicity of infection. ***p<0.001, unpaired two-tailed Student T-test.

**Figure 2.**
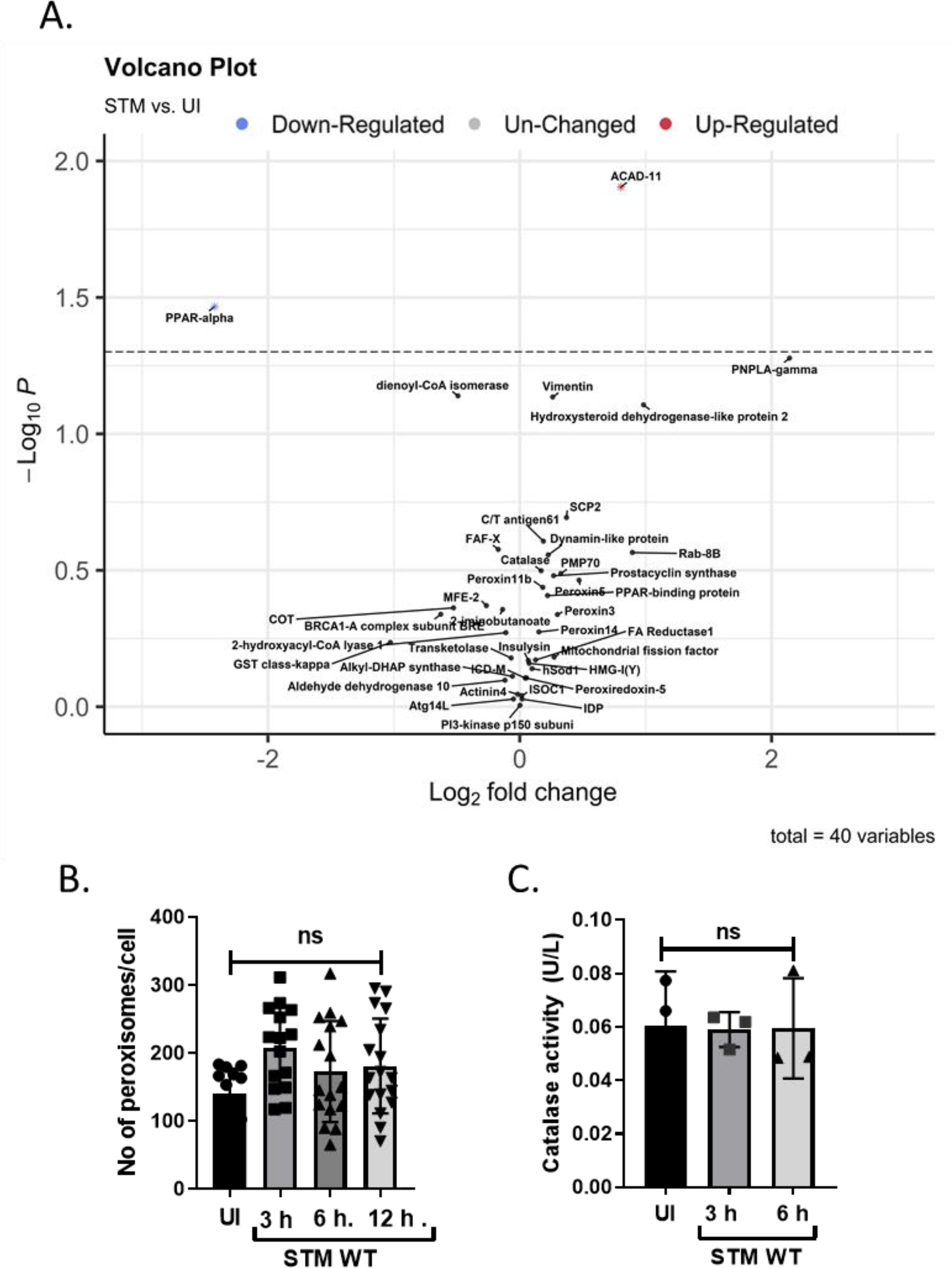
Peroxisomal RNA expression and metabolic pathways are not altered after ***Salmonella*** infection. A. MS-based quantification of peroxisome proteins after 6 hrs of infections in HeLa cells with STM, where, among total 3412 proteins, 40 proteins were associated with peroxisome and shown in the Volcano plot. Samples were compared between uninfected and infected cells. Black dots indicate peroxisomal-associated proteins. B. Schematic graph showing peroxisome number per HeLa cell after STM infection at different time points (3,6 and 12 hours) in comparison to the uninfected cells. In the graph each dot represents one cell (total cells n=20). Similar results were confirmed in three technical replicates; C. Catalase activity in HeLa cell after STM infection with indicated time points. Individual data points represent mean ±SD. Result is representative of 3 independent experiments. ‘MOI’ denotes multiplicity of infection. ‘ns’ denotes non- significant, unpaired two-tailed Student T-test.

### Peroxisomes are required for efficient intracellular replication of ***Salmonella*** Typhimurium

To better understand if this altered temporal dynamics benefit the host by inducing immune response or help the bacteria to survive better, we generated PEX5 knockout cell line using CRISPR/Cas9 in HeLa cells confirmation in (Supplementary figure 3A-C). PEX5 is the major protein that binds to peroxisomal target sequence-1 (PTS1) containing proteins and help them shuttle to peroxisomal matrix. We found that the PEX5 KO cells which are devoid of functional peroxisomes showed reduced *Salmonella* replication (Figure 3A) and lesser number of SCVs per cell (Supplementary figure 3D). Additionally, treatment with 4-PBA chemical inducer of peroxisome biogenesis, heightened PEX14 expression in HeLa cells (Supplementary figure 3E) and displayed enhance bacterial replication (Figure 3B) suggesting that functional peroxisomes are required for intracellular *Salmonella* proliferation. None of the siRNA and inhibitor tested altered cell viability (Supplementary Figure 3F-G).

**Figure 3.**
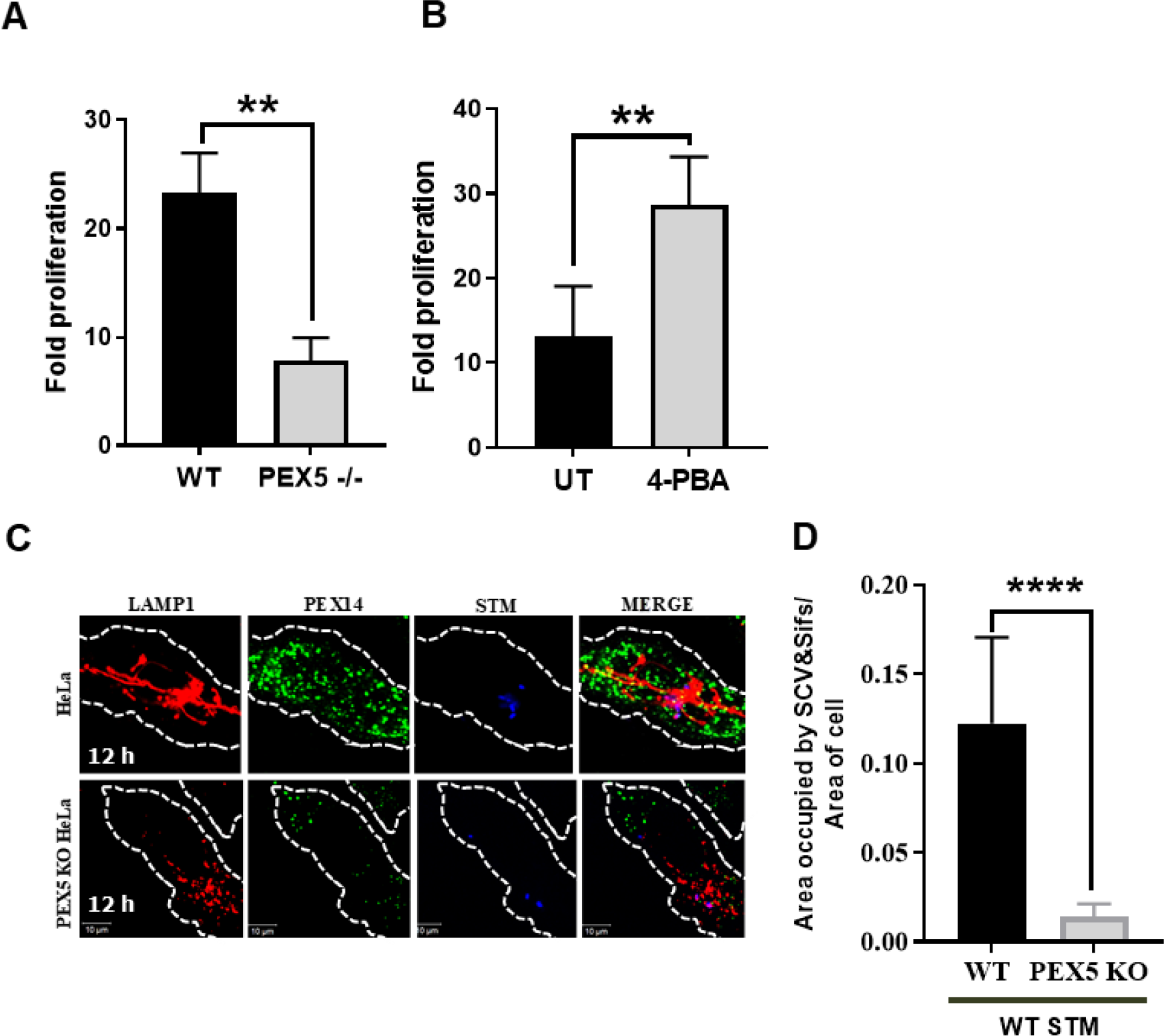
Peroxisomes are required for efficient intracellular replication of ***Salmonella*** Typhimurium. A. STM proliferation in WT and PEX5^-/-^ HeLa cells after 16 hr of infection with STM (MOI=10). CFU was shown as fold change compared to 2 hr. The graph is representative of three independent experiments with similar results. B. HeLa cells were treated with 5 mM 4-PBA for 10 days. Those cells were infected with STM (MOI=10) and CFU was shown as fold change compared to 2 hr. The experiment was repeated 3 times. C. WT and PEX5^-/-^ HeLa cells infected with STM for 12 hours. Cells were fixed and stained with LAMP1 (red), Pex14 (green) and STM (blue). D. Graph represented the area occupied by SCVs with SIFs in WT and PEX5^-/-^ HeLa cells after 12 hours of STM infection (MOI=50). Cell number, n=20; scale bar 10 um. Individual data points represent mean ±SD. ***p<0.001, **p<0.01, unpaired two-tailed Student T-test.

To further verify if peroxisomes contribute in formation of SCVs, infection was performed in the PEX5 KO HeLa cells and formation of filaments post infection was studied temporally. We observed a decreased number of SCVs in PEX5KO HeLa cells (Supplementary figure 3D). Additionally, among the SCVs present, the filament length of SIFs was drastically reduced in case of PEX5 KO HeLa cells as compared to WT HeLa as quantified by area occupied by SCVs containing SIFs (Figure 3C-D). Thus, collectively it can be deduced that peroxisomes help in the formation of filaments that are essential during maturation of SCVs and bacterial growth. These results establish peroxisomes as a pro-*Salmonella* organelle.

### ***In silico*** analysis identified a ***Salmonella*** effector protein, SseI to contain putative peroxisome targeting sequence

Examination of HeLa cells infected with the *ΔssaV* strain (where functional SPI2 needle is absent and bacterial SPI2 effector proteins are not secreted) showed no colocalization with peroxisomes (Figure 4A). We assumed that a bacterial effector protein might be playing a crucial role in the peroxisome-SCV dynamic contacts that we observed in the WT *Salmonella* infected cells and carried out an *in silico* analysis of 127 *Salmonella* T3SS effector proteins to identify proteins that contain putative host peroxisome targeting sequence 1 (PTS1). The analysis predicted SseI, a SPI-2 encoded effector to contain a variant of PTS1 tripeptide, ‘GKM’. The presence of this PTS1 motif in SseI was conserved across multiple *Salmonella* serovars infecting many hosts (Fig.4B). Microscopic analysis of SseI deletion strain exhibited reduced interaction with peroxisomes (Figure 4C-D). The *ΔsseI* strain and the *ΔssaV* strain showed drastic growth reduction in the HeLa cells (Figure 4E). Further, to experimentally verify SseI gets targeted to peroxisomes, we generated N-terminal HA tagged clones containing full length SseI and expressed the clones in HeLa cells. We observed presence of SseI on purified peroxisomes, and specifically on the peroxisomal membrane. Removal of either predicted PTS1 tripeptide or point mutation in PTS1 resulted in reduced SseI accumulation on peroxisomes (Figure 4F-G). The validity of the assay system and purity of peroxisomal isolation is shown in Supplementary figure 4A-B. To characterize the importance of this interaction between SseI and peroxisomes, HeLa cells transfected with these constructs were infected with SseI deletion strain of *Salmonella*. We observed that the *Δ sseI* strain had a drastic growth defect consistent with the reduced peroxisomal colocalization. Further, rescue in bacterial replication was observed in case of cells expressing full length SseI compared to cells expressing the PTS1 mutants (Figure 4H).

**Figure 4:**
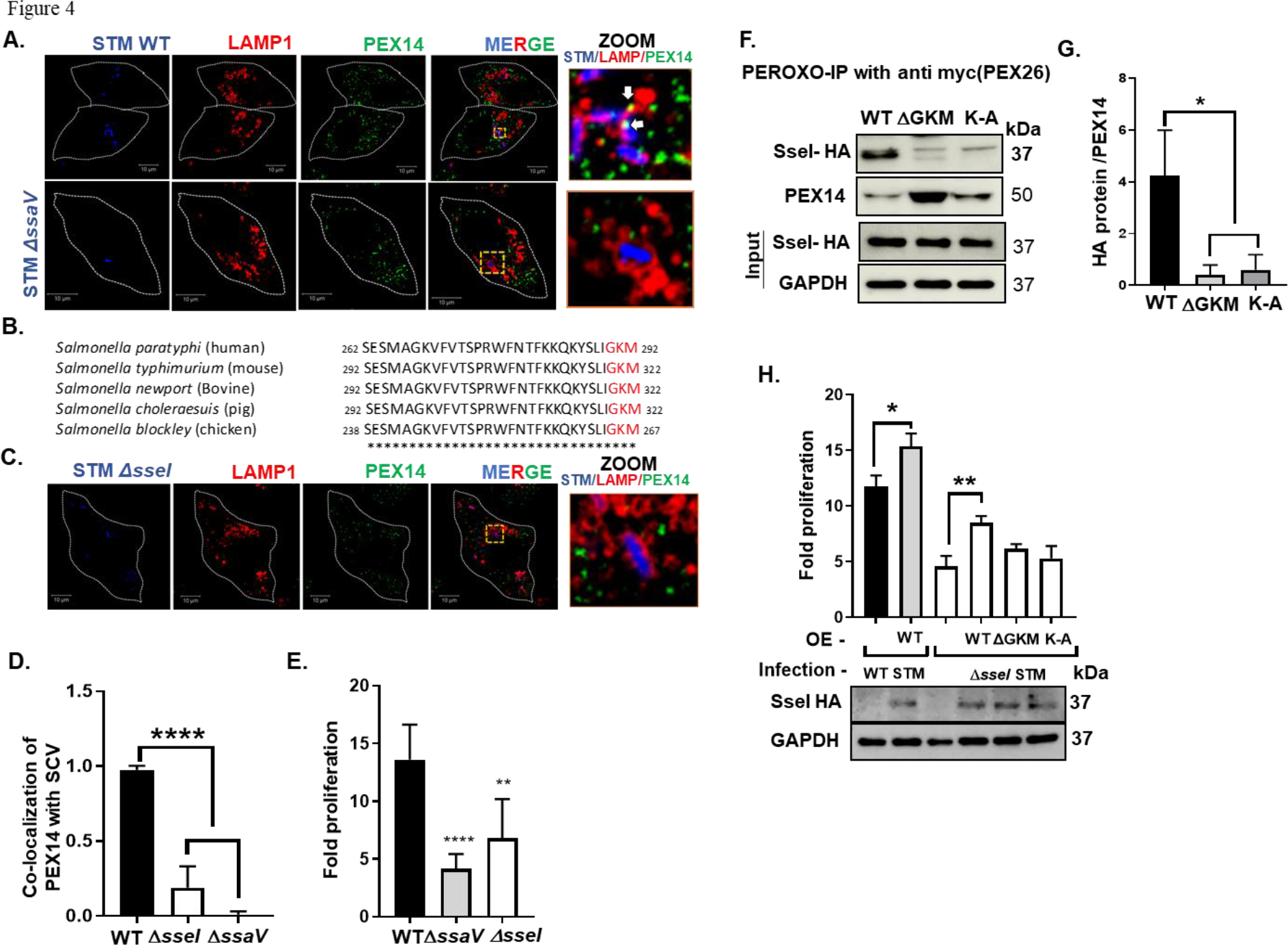
***In silico*** analysis identified a ***Salmonella*** effector protein, SseI to contain putative peroxisome targeting sequence. A. HeLa cells were infected with either the wild type (WT) or *ΔssaV* strain for 12 hours; Cells were fixed; stained for PEX14(green), LAMP1(red), and STM (blue); and Zoom part of image are shown (white arrow) colocalization pex14(green) with SCV(LAMP1); scale bar 10 um. B. Conserved PTS1-like sequence (GKM) in the C-terminal 30 amino acid residues of multiple *Salmonella* species infecting different hosts is depicted. C. Representative microscopy images of HeLa cells infected with the ΔsseI STM for 12 hours; Cells were fixed; stained for PEX14(green), LAMP1(red), and STM (blue); and Zoom part of image were showing pex14(green) with SCV(LAMP1); scale bar 10um. D. Graph representing the percentage co-localization of Pex14 with ‘LAMP1 labelled SCVs’. n=20 cells per group E. Hela cells were infected with either the wild type (WT) or *ΔsseI* or *ΔssaV* strains and fold increase in bacterial growth from 2 h to 12 h is shown. F. Peroxisomes were isolated by PEROXO-IP (ANTI-MYC: PEX26) from HeLa cells transfected with N-term HA tagged SseI clones (full length/PTS1 deletion - ΔGKM/point mutation on PTS1-> K-A) and SseI migration to purified peroxisomes were quantified using western blot. The immunoblot was incubated with anti-HA (for SseI) and Anti-Pex14 Antibody (to check peroxisome purity. G. Quantification for HA: SseI is shown where band intensity was normalized to PEX14. H. Fold proliferation of the wild type (WT) and STM *ΔsseI* strain after transfecting HeLa cells with N-term HA tagged SseI clones (full length/PTS1 deletion - ΔGKM/point mutation on PTS1-> K-A). The transfection efficiency was further validated using western blot and shown below. The experiments were repeated at least two times. ‘MOI’ denotes multiplicity of infection. Individual data points represent mean ±SD., ****p<0.0001, *p<0.05, **p<0.01, unpaired two-tailed Student T-test.

### SseI binds to host GTPase ARF1 to regulate its activity

It was tempting to speculate that SseI might sequester/activate functionality of some host protein on the peroxisome. From the available literature ^37, 38^ we know certain proteins are predicted to interact with SseI (Supplementary figure 5A). One among them, ADP-ribosylation factor 1 (ARF1) is a small GTPase and is involved in protein trafficking. It is further known to interact with PEX35 in the peroxisome and also involved in vesicle migration^39^. Recent studies have indicated a role for ARF1 in synthesis of PIP2, which is essential for organelle docking and movement^31, 40^. Next, conducted the molecular docking of our target proteins (ARF1 and SseI) using HDOCK SERVER, a protein-protein or protein-DNA/RNA interaction platform and observed indeed SseI and ARF1 can interact (Supplementary Figure 5B). Co-immunoprecipitation of GFP-tagged ARF1 and HA-tagged SseI (Figure 5A) demonstrated that these two proteins interact in HeLa cells. To directly show binding, SseI protein was purified (Supplementary Figure 5C) and it showed binding with HA-tagged ARF1 from overexpressed cell lysates (Figure 5B). We next observed that endogenous ARF1 gets recruited on the peroxisomal membrane and this re-localization is enhanced after *Salmonella* infection (Figure 5C). Following interaction, shown in figure 5D, infection with wild type *Salmonella* activates ARF1 more when compared to the *ΔsseI* strain. Thus, these results suggest that SseI mimics the role of a host ARF1 GEF during infection. Following this, we tested the importance of ARF1 activation in *Salmonella* proliferation by silencing ARF-1.

**Figure 5.**
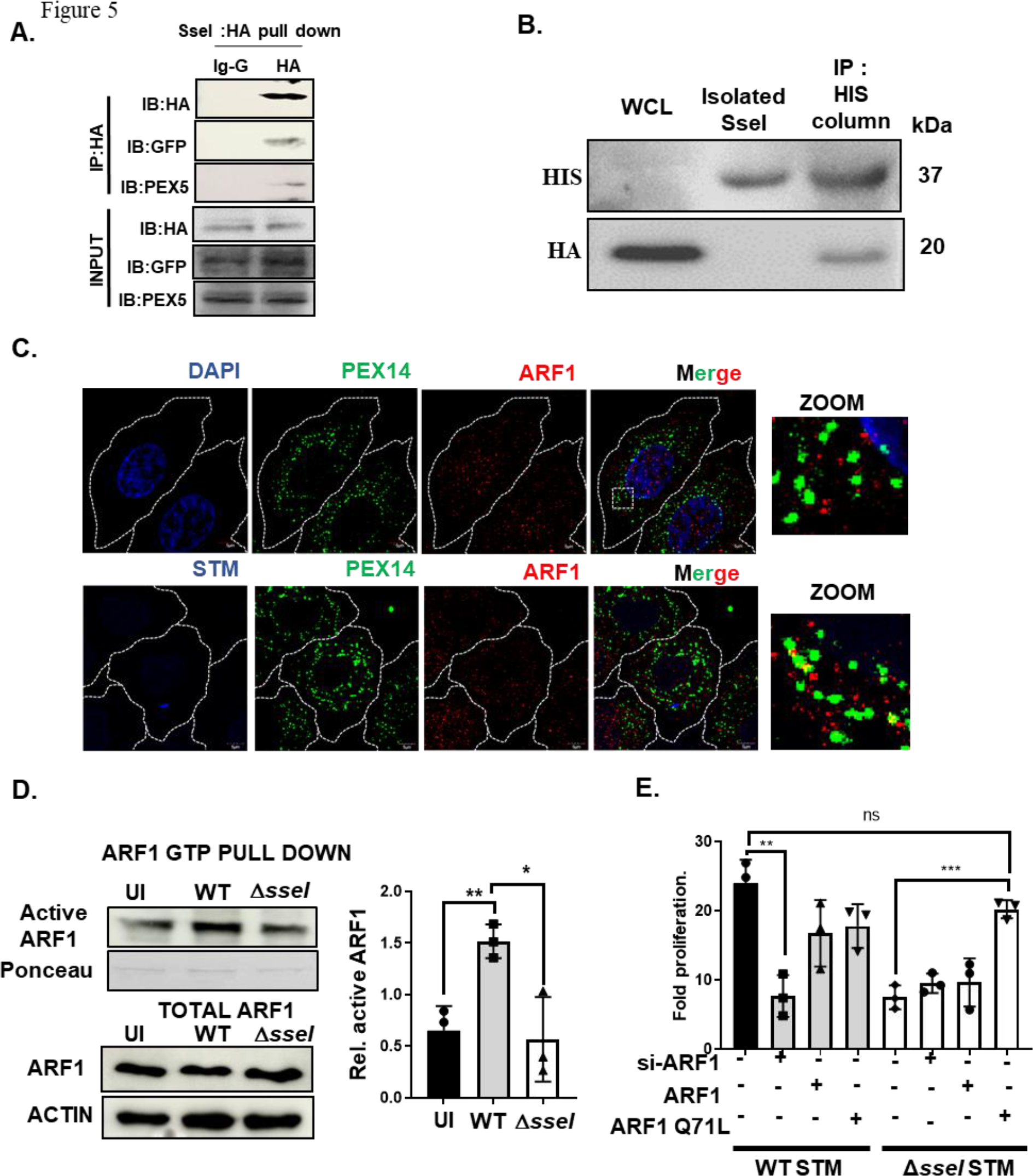
SseI interacts with host GTPase ARF1 on peroxisome. A. HeLa cells were transfected with HA: SseI and GFP: Arf1 plasmids and then immunoprecipitated as described in the experimental procedures. Blot was incubated with anti-HA for SseI, anti-GFP for ARF1 and anti-PEX5. Input indicates the total cell lysate of the cells mentioned above. B. Total cell lysates from HeLa cells were transfected with HA: ARF1 was collected and incubated with purified his tagged SseI for 2 hr. Then his tagged SseI was pulled down by Ni-NTA beads and the eluted sample was run on SDS-PAGE. Immunoblot analysis for His-tagged SseI and HA-tagged ARF1 is shown. Input indicates total cell lysate from the ARF1: HA transfected cells. C. Representative microscopy images of HeLa cells infected with STM (MOI=50) for 6 hours; Cells were fixed; stained for endogenous ARF1(red), Pex14(green), and STM-DAPI (blue); and Zoom part of image are shown ARF1(red), Pex14(green), and STM-DAPI (blue) in both infected and uninfected cells; scale bar=5um. D. Equal number of HeLa Cells were infected with either the WT or STM *ΔsseI* (MOI=10) for 6 hr. Then the active ARF1 was pulled down and were run on SDS page. Densitometry graph from 3 independent experiment representing the relative ARF1 activation in the WT and *ΔsseI* infected cells compared to that of uninfected HeLa cells is shown (MOI=10). E. HeLa cell were transfected with either full length or mutated (Q71L) ARF-1 or silenced ARF-1. Then infected with the WT or *ΔsseI* STM and changes in the fold proliferation were plotted. Experiments were repeated three times. Individual data points represent mean ±SD. Result is representative of 3 independent experiments. ‘MOI’ denotes multiplicity of infection. ‘ns’ denotes non-significant ***p<0.001, *p<0.05, **p<0.01, unpaired two-tailed Student T-test.

Interestingly, we observed a reduction in the bacterial proliferation during ARF1 silencing during WT *Salmonella* infection. On the other hand, in the cells where constitutive active ARF1 (Q71L) was expressed, the growth defect of the *ΔsseI* strain reverted similar to the WT strain (Figure 5E). We could effectively knockdown ARF-1 (Supplementary figure 5D) and ARF1 knockdown had marginal effect on cell viability (Supplementary figure 5E).

### Activated ARF1 induces PIP2 synthesis to strengthen the peroxisome contacts with SCVs

ARF1 localizes to yeast peroxisomes and highly purified rat liver peroxisomes and regulate phosphatidylinsolitol-4-5-bisphosphate (PI-4,5-P2 or PIP2) levels at the membrane in HL60 cells by activating phosphatidylinsolitol-4- phosphate-5 kinases (PI5P4Ks)^31, 41^. Previously PIP2 has been shown to help in organelle contact and also organelle movement^30, 42, 43^. Given, ‘peroxisome contacts with lysosomes and SCV’ increased after infection as well as peroxisome movement, we looked closely at PIP2 at the organelle level. We tested the possibility of *Salmonella* infection leads to an enhanced PIP2 synthesis from the host.

Studies show that activated ARF1 induces PI (4,5) P2 synthesis and we therefore next asked if WT *Salmonella* induces ARF1 activation leading to subsequent PIP2 synthesis in the peroxisome membrane. We observed that WT *Salmonella* infection led to an enhanced ARF1 activation after 6 hours of infection coinciding with the enhanced PIP2 time course (Fig.6A). Notably, infection with the *Δ sseI* strain failed to induce PIP2 accumulation on the peroxisome pointing out the role of SseI in this connection. We also observed PIP2 staining on the peroxisome membrane (Supplementary figure 6B). To rule out if SseI can directly bind to any of the phospholipoid including PIP2, we did protein-lipid overlay assay by using PIP strip membrane, and found it binds to none of the lipids tested (Supplementary figure 6A).

**Figure 6:**
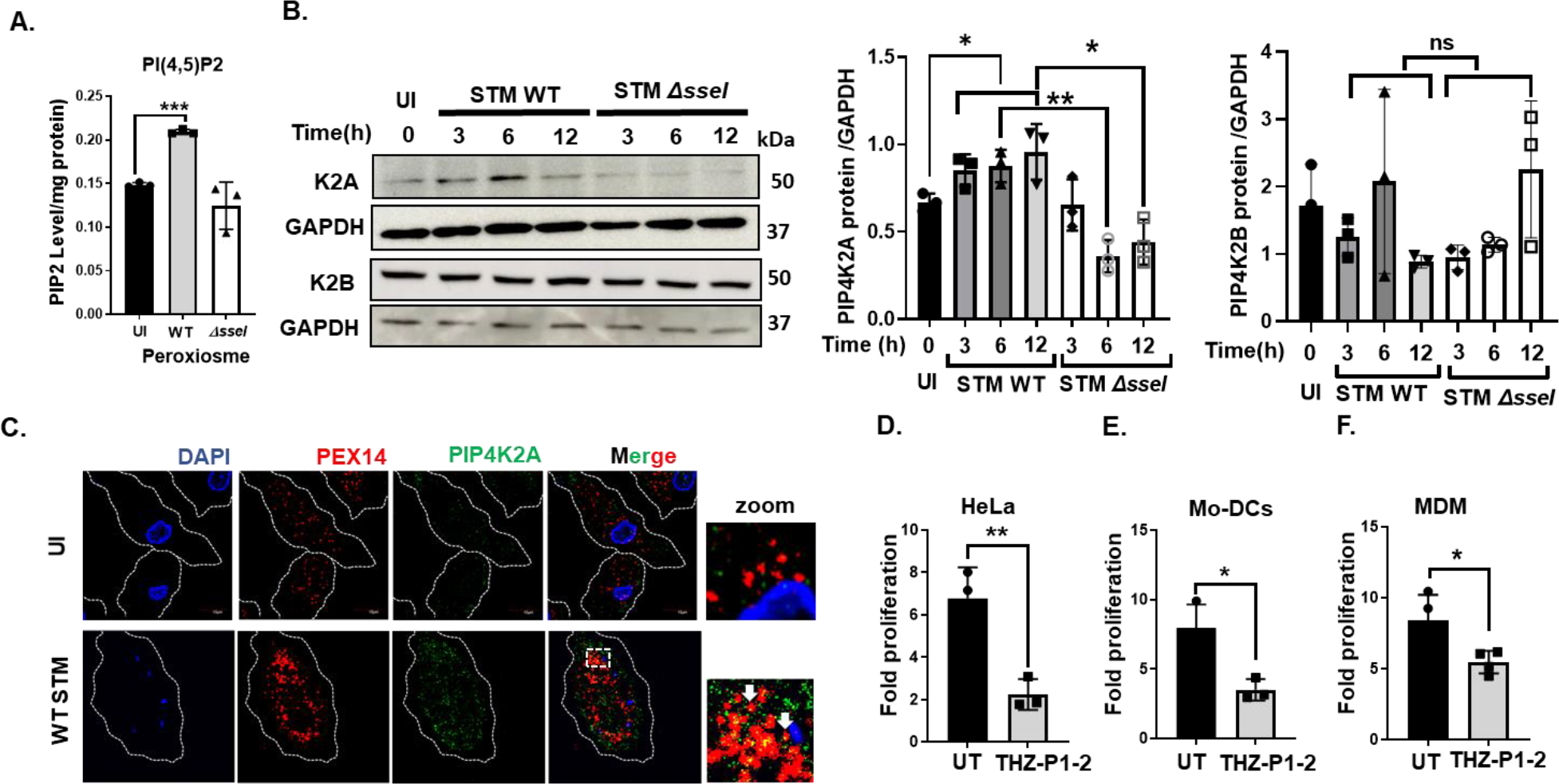
Activated ARF1 induces PIP2 synthesis by recruiting PIP4K2A kinase to the peroxisome A. PI (4,5) P2 level on peroxisome was measured by ELISA in uninfected and infected with the WT or STM *ΔsseI* (MOI=10, for 6h) in the PEROXO-tagged stable cell line where cells were transfected with lentivirus overexpressing 3X myc-EGFP-Pex26. B. Immunoblot analysis of PIP4K2A and PIP4K2B protein level in the cells after wild type (WT) or STM *ΔsseI* infection for 3,6, and 12 hr (MOI=10). ‘UI’ denotes uninfected cells. Densitometry analysis was done from n=3 independent experiments. C. Representative microscopy images of HeLa cells infected with STM for 6 hours; Cells were fixed; stained for PIP4K2A (green), PEX14(red for peroxisome), and STM (blue); and Zoom part of image are shown. PIP4K2A (green), PEX14(red for peroxisome) colocalization in both infected and uninfected cells; (n=20 cells, and similar results wereobserved in independent experiments). Scale bar 10 um. D. HeLa Cells (E) Human monocyte-derived dendritic cells (Mo-DCs) (F) Human monocyte-derived macrophages (MDM) were treated with PIP4K inhibitor (THZ-P1-2, 1 uM) and infected with STM (MOI=10). Fold change of STM after 16 h was quantified by counting colony forming units (CFU) and compared between treated and untreated HeLa cells. ‘UT’ denotes untreated cells. Result is representative of 3 similar experiments. ‘MOI’ denotes multiplicity of infection. Individual data points represent mean±SD ‘ns’ denotes non- significant ***p<0.001, *p<0.05, **p<0.01, unpaired two-tailed Student T-test.

We next wished to evaluate the regulators of PIP2 synthesis downstream of active ARF1, specifically in the peroxisome membrane. Although PI4P5K is responsible for the synthesis of most of the cellular PIP2^44^, the non-canonical phosphatidylinsolitol-5-phosphate-4 kinases (PI5P4K/PIP4K) ^45^regulate membrane PIP2 generation^46^, however remains understudied during infection. Multiple PIP4K isoforms exists in the cell and we observed that *Salmonella* infection specifically upregulate PIP4K2A, not the PIP4K2B isoform during infection (Fig. 6B).

Recruitment of PIP4K2A to the peroxisome was next assessed and indeed Salmonella infection led to a higher peroxisome targeting of this isoform (Fig. 6C). Blocking this kinase with a specific inhibitor THZ-P1-2^47^ leads to a drastic growth defect of the WT strain in Hela cells and in human blood derived dendritic cells (Mo- DC) and human blood derived primary macrophages (MDM) and (Fig. 6D-F). THZ- P1-2 did not change cell viability at the concentration tested (Supplementary Fig.

6C). Altogether, our data till now clearly proved that SseI is recruited to the peroxisome, binds and activates host ARF1 on the peroxisomes leading to an increase in the levels of PI (4,5) p2 on peroxisomes.

### LDL-derived cholesterol routed via lysosome-peroxisome is essential for ***Salmonella*** growth

Why will an intracellular bacteria display such an elaborate mechanism to alter peroxisomal PIP2 level to enhance peroxisome moment and/or connection with SCV? Published literature depicted that Lysosomal Syt7 binds to PIP2 and mediate cholesterol transport from lysosome to peroxisomes^24^. And, PIP2 generated at the peroxisome is essential for lipid trafficking to mitochondria and ER^45^. As infection with the SSE- mutants leads to reduced bacterial load and SIF formation with a reduced PIP2 level, we tested the possibility of *Salmonella* infection leading to a higher peroxisomal-lysosomal contacts. Indeed, there was enhanced lysosome to peroxisome contacts during *Salmonella* infection (Fig.7A) and previous studies reported cholesterol to be essential in SCV maturation.

**Figure 7:**
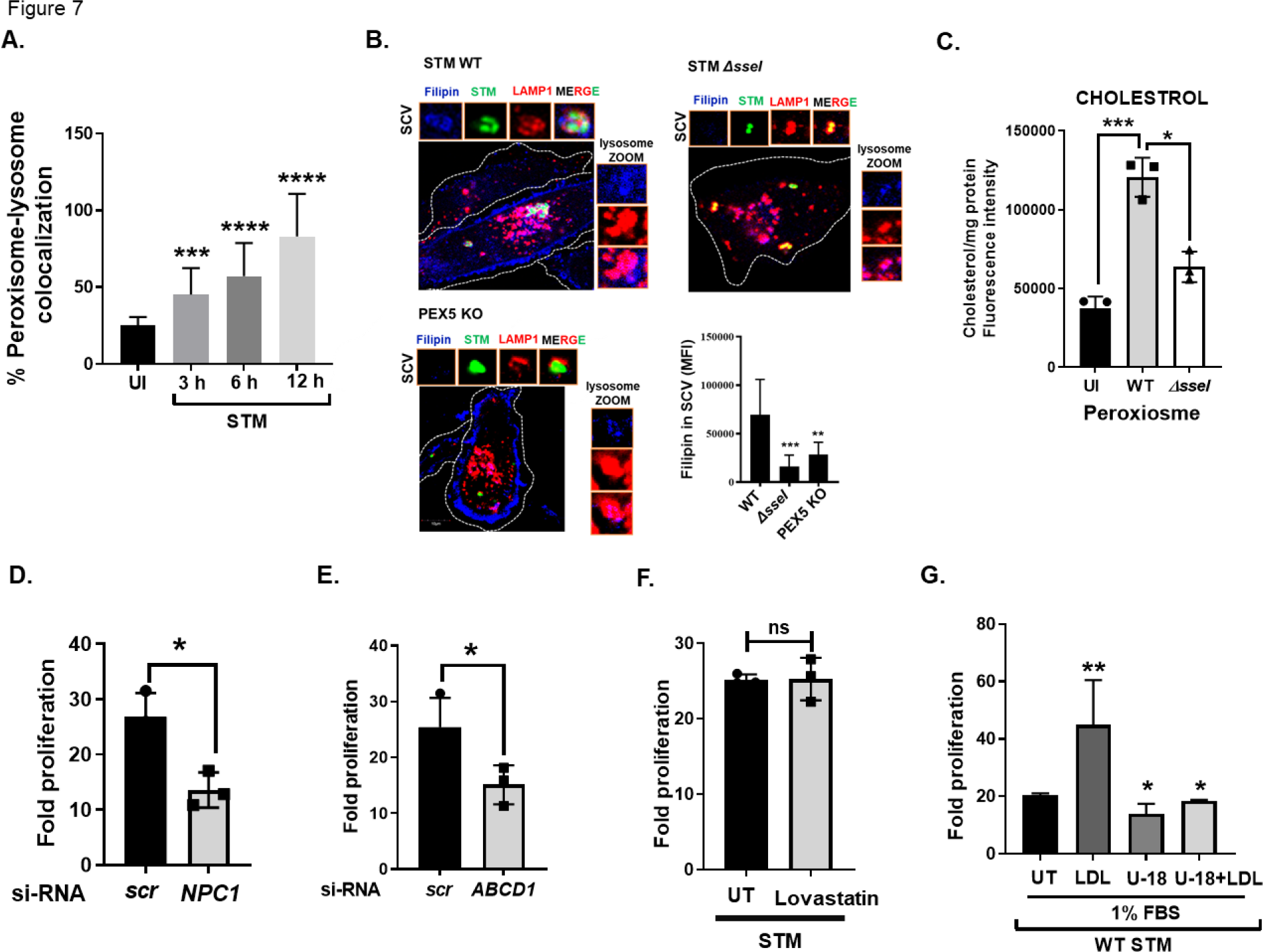
LDL-derived cholesterol routed via lysosome-peroxisome is essential for ***Salmonella*** growth A. Graph representing percentage of peroxisome-lysosome co-localization in the HeLa cells after infection with the WT STM (MOI=10) for different time points. B. WT HeLa cells were infected with WT STM or STM *ΔsseI* for 6 h with MOI= 50. pex5^-/-^ HeLa cell were infected with WT STM for 6 h with MOI= 50. Cells were fixed and immunostained with LAMP1(red), cholesterol using filipin(blue) and STM (green). Zoom part represented SCV and Lysosome cholesterol (blue) level in the cells. Graph representing change in the filipin fluorescent intensity is shown. (n=50 cells). Scale bar 10 uM. C. PEROXO- tagged stable cell was generated using transfection with lentivirus overexpressing 3X myc-EGFP-Pex26. Stable PEROXO tag HeLa cells were next infected with the WT and *ΔsseI* STM with MOI=10 for 6 hr. Isolated peroxisomal cholesterol amount was measured using cholesterol assay kit. Graph representing the fold proliferation of STM silencing D. NPC1 E. ABCD1 in HeLa cells. Changes in fold proliferation of WT STM after 2 and 16 hr post-infection is plotted in these cells. F. Cells were kept untreated or treated with lovastatin (10uM). Changes in fold proliferation of WT STM after 2 and 16 hr post-infection is plotted in these cells. G. Cells were kept untreated or treated with LDL/U18666A in the media containing 1% FBS in HeLa cells. Changes in fold proliferation of WT STM after 2 and 16 hr post-infection is plotted in these cells. Individual data points represent mean± SD. Result is representative of 3 independent experiments. ‘MOI’ denotes multiplicity of infection. ‘src’ denoted scrambled siRNA. ‘UI’ denotes uninfected, ‘UT’ denotes untreated. ‘ns’ denotes non- significant ****p<0.0001, ***p<0.001, *p<0.05, **p<0.01, unpaired two-tailed Student T-test.

Hence, we focused our attention if the *Salmonella* induced enhanced cholesterol transfer to peroxisome later gets conveyed to the SCV as direct binding to lysosome is detrimental for bacterial growth. In order to test this, we stained cellular cholesterol using fluorescent dye filipin that specifically binds to cholesterol and we found that WT *Salmonella* induces cholesterol accumulation in the SCV. The *Δ sseI* strain failed to sequester cholesterol in the SCV and interestingly in that scenario the cellular cholesterol accumulated in the lysosome. In the PEX5 mutant HeLa cells, the WT *Salmonella* infection mimics the *Δ sseI* phenotype like the WT Hela cells confirming that peroxisome serves as the bridge between lysosome and SCV during cholesterol movement (Fig. 7B). We validated cholesterol accumulation on isolated peroxisomes and data shown in Fig. 7C indicates higher cholesterol content when infected with the wild type bacteria after 6 hours of infection. In case of *Δ sseI* strain there was reduced cholesterol in that condition. Taken together, these results demonstrate that LDL-derived cholesterol in lysosomes accumulates in SCVs during maturation.

To understand this better, we next silenced either the lysosomal cholesterol transporter, NPC1 or peroxisome membrane protein, ABCD1 whose expression is regulated by cholesterol levels (Supplementary figure 7A-B). Knockdown of these proteins are known to perturb the lysosome-peroxisome contacts thereby reducing cholesterol transport to peroxisomes. As seen in figure 7D-E, silencing of either NPC1 or ABCD1 showed reduced intracellular *Salmonella* replication. Treatment with inhibitor of de novo cholesterol synthesis, lovastatin did not show any perturbation in the rate of bacterial replication (Figure 7F) making us believe that indeed *Salmonella* require exogenous cholesterol for growth. Next set of experiments supported this idea. Exogenous addition of LDL to media enhanced the bacterial replication and addition of a NPC1 chemical inhibitor, U18666A either alone or along with LDL prevented bacterial replication (Figure 7G). None of the siRNA or inhibitors tested altered cell viability (Supplementary Fig.7C-F).

SYT7 on SCV could help in the peroxisome docking and cholesterol transfer.

Transport of molecules between organelles requires participation of tethering proteins to bring the organelles in close proximity thereby facilitating transfer. We next wanted to identify the binding partner on SCVs that interact with peroxisomes. NCP1, synaptogamin VII (Syt7) and extended synaptogamin (E-Syt) present on lysosomes and ER respectively were known primarily interact with peroxisomal PI(4,5)P2 to transport cholesterol^24, 30, 48^. Next, peroxisomal cholesterol accumulation was measured in the NPC1, SYT7 and Arf-1 knockdown cells and a significant reduction of cholesterol in the purified peroxisomes was observed (Fig.8A). To improve our understanding of the mechanism, we examined the presence of Syt7 and E-Syt on SCVs. Among the two proteins tested, Syt7 showed localization on SCVs (Figure 8B) and maintains SCV cholesterol level. In addition, when cells were silenced with Syt7 (Fig.8C), bacterial replication was restricted (Fig.8D). In contrast, E-Syt neither showed localization with SCVs nor silencing caused any change to bacterial proliferation (Supplementary figure 8A-C). None of the siRNA tested altered cell viability (Supplementary figure 8D-E). This data demonstrates that LDL-derived cholesterol in lysosomes gets transported to SCVs use peroxisome as a bridge. Altogether, these experiments suggest that transport of cholesterol from peroxisomal to SCVs is mediated by PI (4,5) P2 on peroxisome membrane and Syt7 on membrane of SCVs.

**Figure 8:**
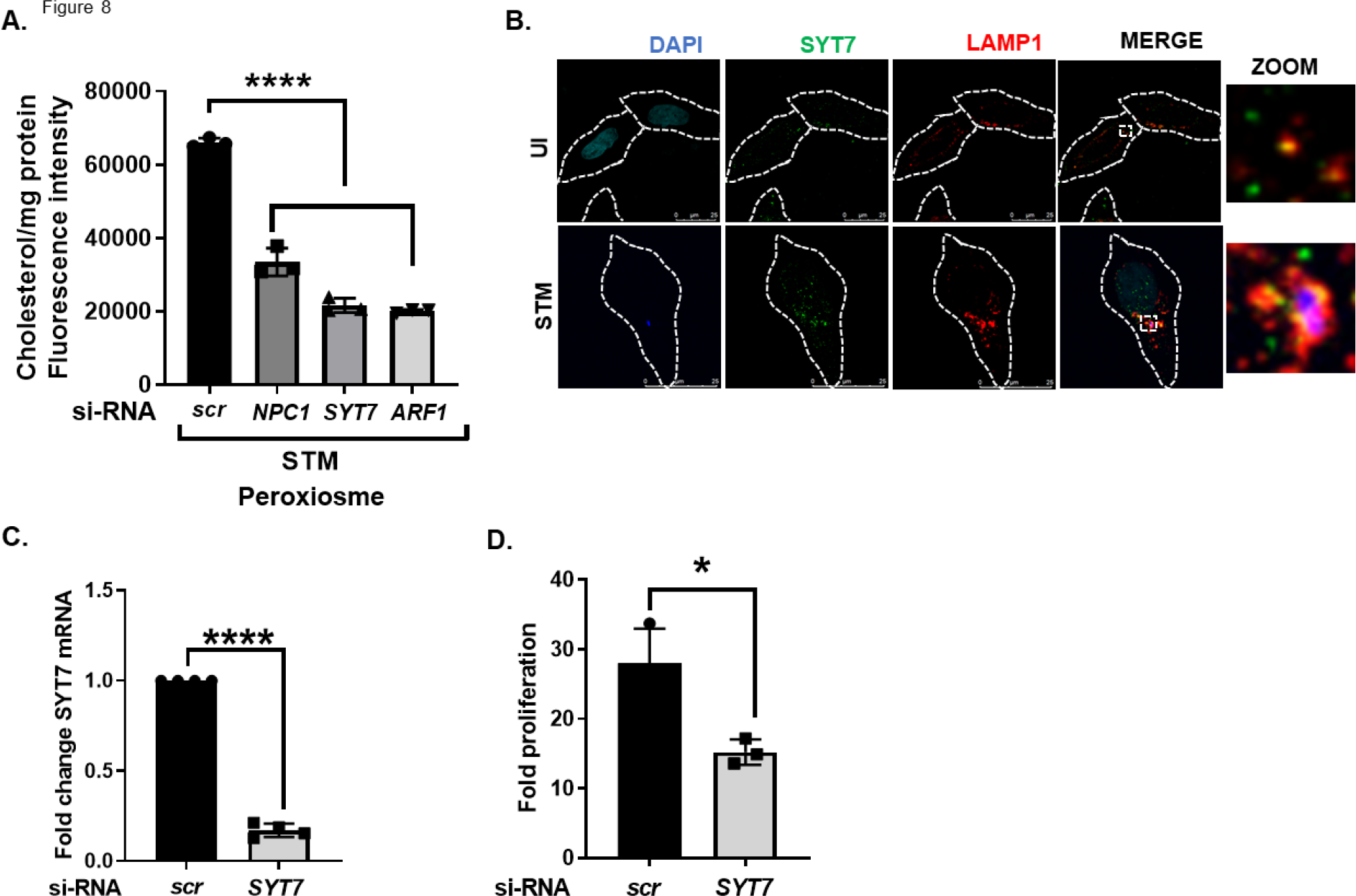
SYT-7 present on SCV is essential for interaction with peroxisome. A. PEROXO-tagged stable cell was generated using transfection with lentivirus overexpressing 3X myc-EGFP-Pex26. Stable PEROXO tag HeLa cells were next transfected with non-targeting si RNA (scr) or si RNA against NPC1, SYT7 and ARF1 and incubated for 48 hours. Then the transfected cells were infected with WT STM with MOI=10 for 6 hr. Peroxisomal cholesterol is shown as measured by using cholesterol assay kit. B. Representative microscopy images showing syt7(green) co-localization with lysosome (LAMP1) and SCV (red) with STM of MOI=50, scale bar 25 uM. C. HeLa cells were transfected with non-targeting si RNA (scr) or si RNA against syt7 for 48 hr. Silencing efficiency of syt7 was measured by q-PCR. D. Changes in fold proliferation of WT STM after 2 and 16 hr post-infection is plotted in these cells. Individual data points represent mean ± SD. Result is representative of 3 independent experiments. ‘MOI’ denotes multiplicity of infection. ‘ns’ denotes non- significant ****p<0.0001, *p<0.05, unpaired two-tailed Student T-test.

### SseI mutant is attenuated in the animal model of infection

To gain further evidence, whether SseI is essential for *Salmonella* pathogenicity, we employed a mouse model of infection and orally inoculated the animals with the WT or the *Δ sseI* strain. Data shown in Fig. clearly indicates that after 7 days’ post-infection, the mutant strain grew significantly less when compared to the wild type counterpart. This hypo virulence in the *Δ sseI* strain was observed in all the mice organs like MLN, Intestine, liver and spleen (Fig. 9) indicating the in vivo essentiality of SseI and the peroxisomal pathways involved in the mechanistic pathways identified before.

**Figure 9:**
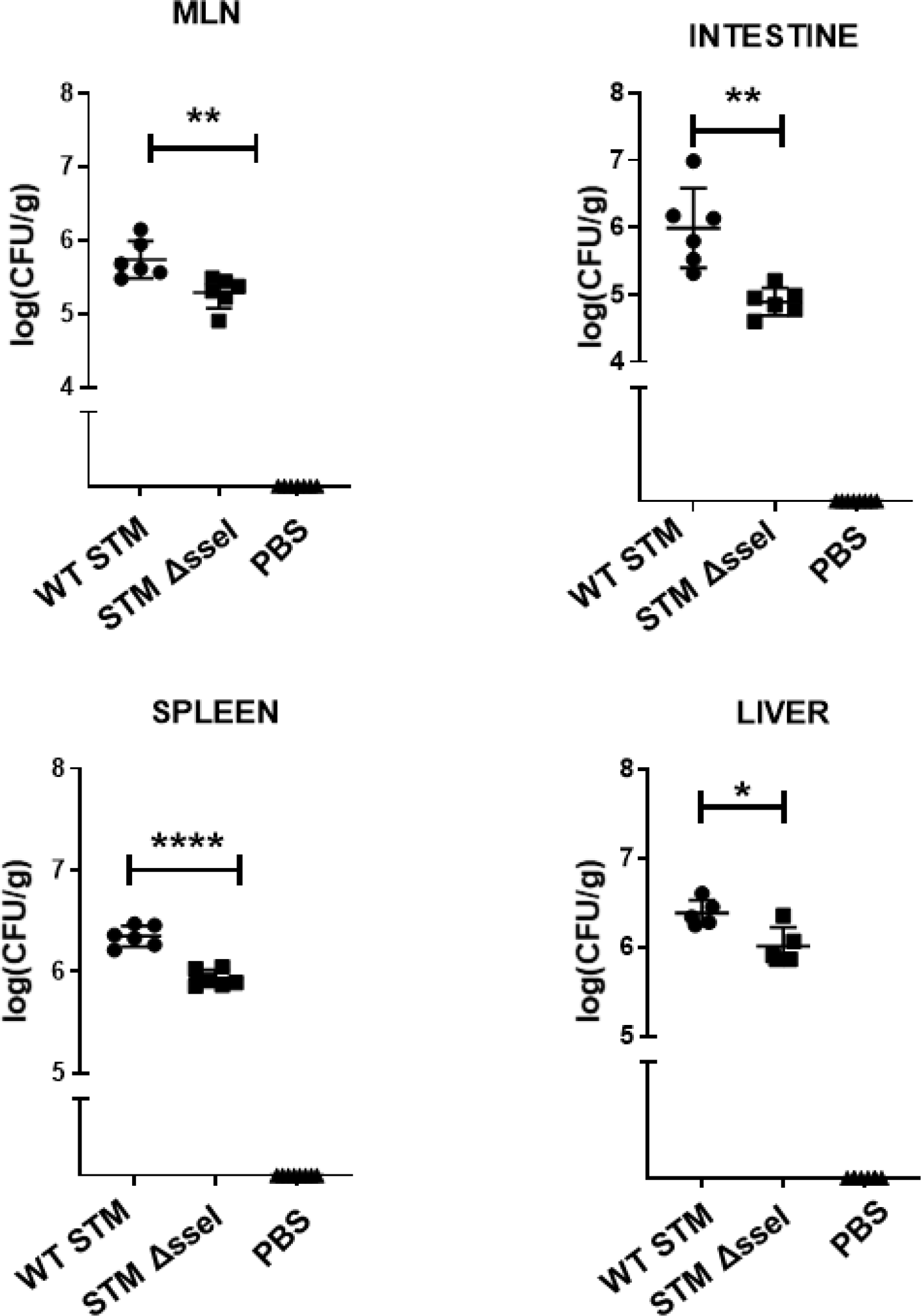
*ΔsseI* is attenuated in the animal model of infection. 5-6 weeks old C57BL/6 mice were infected with 10^7^ CFU of the WT or STM *ΔsseI* strain via oral route, PBS was used as a vehicle control (n=6). Mice were sacrificed after 7 days and *Salmonella* burden was enumerated by plating. CFU/g of each organ (intestine, MLN, Spleen and Liver) was determined. Individual data points represent mean ±SD ****p<0.0001, *p<0.05, **p<0.01, unpaired two-tailed Student T-test.

Collectively, our findings demonstrate that *Salmonella* infection perturbs mammalian peroxisome dynamics by transporting lysosomal LDL derived cholesterol to SCV using peroxisome as a bridge (Supplementary Fig.9). Mechanistically, we establish how a bacterial virulence protein targets host peroxisome and modulate host organellar lipid synthesis.

## Discussion

The severity of an infection depends on the arms race between host susceptibility and pathogen virulence. During viral infection, peroxisomes are morphologically and functionally remodeled and can decide the outcome of infection^49, 50^. On the other hand, role of peroxisomes during bacterial infection in the mammalian cells remains understudied, except during *Chlamydia* infection, where peroxisomes were found to be not essential for bacterial infection^51^. One group has recently shown that peroxisomes are required for phagocytosis of *Escherichia coli* in murine macrophages and that peroxisomes impact cytokine release^22, 23^. In this study, we demonstrate that *Salmonella* infection perturbs peroxisome dynamics to aid its intracellular replication. Infection-induced transient interactions are formed between SCVs and peroxisomes. This interaction enhances the cholesterol transport to SCVs from lysosomes via peroxisomes (Supplementary Fig.9). We observed this phenotype in multiple cell types including epithelial (primary human IEC and HeLa) and immune cells (human primary macrophages and THP1 monocytes).

During infection, pathogens are known to interact with various cellular organelles to modulate their activity and *Salmonella* is no exception. *Salmonella* protein SopA gets recruited to the mitochondria and SSE-G to the Golgi during infection^52, 53^. One of the benefit pathogens achieves by interacting with host organelles is to obtain molecules essential for their replication. Peroxisomes are remarkably plastic concerning their size, number and intracellular localization^54^. All eukaryotic peroxisomal matrix protein contains a tripeptide (prototype SKL) at the C terminus. Parasites like trypanosomes and multiple fungal species have PTS1 containing proteins which are essential for their virulence^55, 56^. Additionally, rotavirus structural protein VP4 contains a PTS-1 sequence and was shown to migrate to host peroxisome^57^. Here, we have identified for the first time a bacterial effector protein that contains functional human PTS1 sequence. *Salmonella* SPI2 effector protein SseI (also called SrfH) had conserved GKM tripeptide at the C terminus and we found that it is also targeted to host peroxisomes in addition to its other cellular distribution. Interestingly, C terminal tagged SseI was previously shown to be targeted to the host cell plasma membrane after palmotylation^6, 58^. We believe that the C-Terminal tag might lead to the loss of functional PTS-1 in SseI. Never the less we also observed a cell membrane localization pattern of SseI. Previous studies have shown SseI to inhibit dendritic cell migration and chemotaxis^10, 59^. On the other hand, SseI promoted murine macrophage motility^60^. Intraperitoneal infection with *ΔsseI* strain in mice have shown ambiguous results. At early time points of infection the *ΔsseI* strain showed enhanced dissemination and at a later time the mutant showed reduced growth in vivo^12, 61^. We perform oral inoculation and observed that *ΔsseI* is attenuated in vivo in all the organs tested after 7 days of infection showing that this is a virulence protein required for efficient in vivo colonization. In the cell lines also the mutant strain showed a prominent growth defect when compared to the wild type *Salmonella*. Mechanistically, WT *Salmonella* showed colocalization with host peroxisome starting from 3 hrs of infection and the *Δ sseI* strain failed to show this interaction making us hypothesize that SCV-peroxisomal contacts are required for efficient bacterial growth. In PEX5 knockout HeLa cells growth of WT *Salmonella* reduced significantly again indicating peroxisome as a pro-*Salmonella* organelle.

Further, we identified that SseI binds and activates host GTPase ARF1 to assist in this interaction. Our pull-down and ELISA data demonstrated that activated ARF1 enhances PIP2 levels on peroxisomes. Arf proteins belong to the Ras superfamily of small GTPases that are known to act as regulators of membrane trafficking^62^. Like any other small GTPases, Arf family proteins also undergo cycling between GTP-bound (active) and GDP-bound (inactive) states^63^. Multiple GTPase activating proteins (GAPs) and Guanine exchange factors (GEFs) have been identified to carry out this Arf cycling for specific cellular functions. It is known that certain GAPs of ARF1 (ASAP1) and ARF6 (ACAP1 and ADAP1) are recruited to plasma membrane at the positions of *Salmonella* entry^64^ and *Salmonella* effector SopF binds to ARF1^65^. Depleting these GAPs from *Salmonella*-induced membrane ruffles is shown to reduce bacterial invasion^66^. Here, we show another mechanism by which intracellular Arf is exploited by *S Typhimurium*. We observed that SseI functionally mimics as GEF for ARF1 causing GTP binding and subsequent activation for enhanced peroxisomal PIP2 generation which can modulate the peroxisomal movement and interaction with another organelle including SCV.

Cholesterol serves as the central lipid of mammalian cells that are involved in various functions including signal transduction, steroidogenesis, bile acid synthesis and regulating biological membrane properties (such as curvature, fluidity and membrane fusion)^67^. Additionally, cholesterol plays a critical role during infection^67^. For example, human immunodeficiency virus (HIV) and influenza virus^68^ utilize cholesterol-rich regions on the plasma membrane to facilitate their entry. Further, bacterial pathogens including *Mycobacterium* employ host cholesterol catabolism to acquire nutrients for their survival^69^. Low-density lipoprotein (LDL) circulated in the blood binds to LDL receptors present on the plasma membrane of many cell types and gets internalized^70^. This internalized lipoprotein then gets processed in lysosomes and cholesterol is released. Part of the LDL-derived cholesterol from lysosomes gets transported to peroxisomes. This transfer is mediated by interaction of lipid phosphatidylinositol 4, 5-bisphosphate (PI(4,5)P2) on peroxisomes and lysosomal Synaptotagmin VII protein ^24^. It is already known that ER acquires lysosomal cholesterol by interacting with peroxisomes^30^. Previous studies have also indicated that SCVs are rich in cholesterol^13, 14^. However, SCVs do not fuse with lysosomes until late time periods after infection^37^. Therefore, it was not clear how cholesterol is directed to the bacterial containing vacuoles. Here, we show that in the absence of functional peroxisomes, cholesterol accumulates in lysosomes without reaching to SCVs.

Infection leads to an increase in lysosome-peroxisome and peroxisome-SCV contacts thereby transporting cholesterol. Knockdown of lysosomal cholesterol transporters affect the bacterial replication by perturbing the cholesterol transfer to SCVs needed for its maturation and survival. In the current study, we show that SCVs acquire the cholesterol by recruiting SseI to the peroxisomes, which enhances their interaction with peroxisomes. In the absence of functional peroxisomes, SCVs do not form membranous extensions called SIF which are essential for its stability and survival. Our results are in line with the previous hypothesis that SPI2 is needed for cholesterol accumulation in SCV^13^ and we also provide the compelling mechanism supporting this. This reduction in cholesterol transport in the *Δ sseI* prevents the formation of SIFs thereby causing immature SCVs. Consequently, there is reduced replication of intracellular *Salmonella*.

Growing evidence has highlighted the importance of host-targeted therapeutics that could be developed against numerous related bacteria and our study has long standing implication in this regard. Inflammatory bowel disease patients have higher peroxisome content in the intestinal mucosa^71^ and the patients cannot efficiently clear pathogens including *Salmonella*^72^, supporting our findings. Studies have also shown that patients with *S*. Typhi infection exhibit high cholesterol level. Further, LDL receptor knockout animal are resistant to *Salmonella* infection. In addition, human genetic variation in VAC14, which leads to altered cholesterol metabolism influences susceptibility to *S.* Typhi infection. Specifically, in an interesting study, reduction in VAC14 expression led to enhanced cholesterol in the cell membrane facilitating *Salmonella* invasion^16^. Additionally, by employing cholesterol-depleting drugs such as ezetimibe enhances clearance of *S.* Typhi in zebrafish ^16^. Here, we provide evidence that cholesterol is essential during *Salmonella* proliferation in macrophages and epithelial cells as well and SseI targeted peroxisomes play a crucial role in providing the same to growing SCV. Finally, we observed that the levels of organellar lipid PI(4,5)P2 are regulated by specific phosphatidylinositol 4- phosphate (PIP) 5-kinase (PIP4K) and their inhibition in vitro reduced *Salmonella* proliferation. We speculate that PIP4 inhibition could be a new host targeted therapy in Gastroenteritis and Typhoid fever.

## Methods

### Chemicals and antibodies

Chemicals and media used were as follows: 4-PBA (P21005 from sigma), DMEM (REF. 12800-058 and gibco), RPMI Medium 1640(REF. 31800-014), Ampicillin and Kanamycin (BC0286 - BR BIOCHEM),, Penicillin and streptomycin (A001 and HIMEDIA.), Bovine Serum Albumin (GLR INNOVATIONS – GLR09.020731), RIPA

Buffer (HIMEDIA – TCL131-100ml), Protease inhibitor cocktail (PI) (HIMEDIA- ML051-1ml), Tween-20 (RC1226 – G Biosciences) Triton X-100 (64518 - SRL), Collagenase H (Sigma - 33278623), IPTG (SIGMA - I67581G)and Trypsin EDTA (TCL070 and HIMEDIA) were purchased from HIMEDIA.

Antibodies used (Abs) used were as follows: anti-LAMP1 antibody (D401S from CST), PEX14 Antibody (A7336), ABCD1 (60153-1-Ig), GAPDH Antibody (T0004),

ESYT1 antibody (A15410), anti-HA tag Antibody(2367), Anti-FLAG Antibody(8146 from CST), anti-myc (2276 from CST) anti-pex5 antibody (12545-1-AP from proteintech), ARF1 Antibody (10790-1-AP), Goat anti-mouse IgG Alexa fluor 594(A- 11032), Goat anti-Rabbit IgG Alexa Fluor 488(A-11034) IgG Alexa fluor 594(A- 11032) SYT7 antibody(NBP2-22420), PIP2 ANTIBODY(NBP2-76433), Cy5 goat anti-rabbit IgG (A10523 - Invitrogen), Control Mouse immunoglobulin (Cat#500-M00- 1mg - PEPROTECH), Control rabbit immunoglobulin (Cat#500-P00-500 ug - PEPROTECH) anti-rabbit IgG antibody and horseradish peroxidase (7074P2) were purchased from Cell Signalling Technology. Ni-NTA Agarose (QIAGEN), Protein A Agarose (Invitrogen) and Filipin (SAE0087 from Sigma) were used.

### siRNA and plasmid transfection

HeLa cells were plated on 12-well plate containing glass coverslips. Cells were then transfected with plasmids (9021,60360,139059 - Addgene) using the Lipofectamine 3000 DNA Transfection Reagent for 48 h, according to the manufacturer’s instructions. All siRNAs SMART pool were obtained from Santa Cruz and Dharmacon: siRNA constructs (SC-105086(ARF1); Synaptotagmin VII siRNA (h): sc-41320; E-Syt1 siRNA (h): sc-95714; NPC1 siRNA (h): sc-41588); Pex14 (h): L-012676-00-0005, ON-Target plus^TM^ Control Pool, D-001810-10-20 (Dharmacon^DM^); ABCD (h) sc-41143; non-specific non-targeting pool using Lipofectamine RNAiMAX reagent according to the manufacturer’s instructions. Protein knockdown efficiency was assessed by q-PCR.

### Cell culture

For isolating and culturing, HeLa, THP1 and human primary cells were grown in DMEM (GIBCO) supplemented with 10% fatal bovine serum (GIBCO) along with antibiotic (1% penicillin/streptomycin) and were maintained in 5% CO2 at 37°C.

### Isolation of PBMCs and MDM Human IEC

Buffy coats (≈30 ml) from healthy donors were obtained from the blood bank at the Transfusion Medicine - King George’s Medical University, Lucknow (UP) India. The PBMCs were isolated as follows: first it was diluted 1:2 with 1X-PBS. Then, PBMCs were isolated using density gradient centrifugation using Histopaque-1077 gradient (Himedia, India). Centrifugation was performed at room temperature 400 *g* for

30 min. After isolation, cells were washed two times in phosphate-buffered saline (PBS) then resuspended in RPMI-1640 (GIBCO) media containing antibiotic without FBS. Then, the cells (around 2 × 10^7^ to 3 × 10^7^ PBMCs/ml in 20 ml) were seeded, and allowed to adhere in a 5% CO2 container at 37° for 2 hrs. Adherent cells were then cultured with complete maturation media (RPMI-1640 with 10% FBS, penicillin/streptomycin, 10 ng/ml macrophage colony-stimulating factor (M-CSF) (Genscript) for 5 days for monocyte-derived macrophages (MDM) differentiation.

Media were changed every 2–3 days.

### Transfection and Infection

Cells were seeded on coverslips and were allowed to attach. After overnight incubation, cells were transfected with plasmid constructs using lipofectamine 3000 (Invitrogen) as per the manufacturer’s instructions. After 48 hours post transfection, STM infection was carried out for indicated time points.

### Immunofluorescence microscopy

For staining, cells were first fixed using 2% paraformaldehyde for 15 mins at 37°C. For antibody staining, permeabilization with Triton X-100 (0.3%)/1% BSA in 1X-PBS was followed by primary antibody (1:100) and incubated overnight at 4°C. For filipin (cholesterol) 10% FBS/PBS,1% triton X-100, 50 μg/ml filipin for 1h and primary antibodies (1:100) all together were incubated for 2 hrs at room temperature. Following secondary antibody incubation was done for 1 hour at room temperature. The cover slips were mounted with 50% glycerol. Finally, the cells images were captured using Intravital imaging facility OLYMPUS BX61-FV1200-MPE

### Lentivirus production

Guide RNAs targeting human PEX5 and scrambled were cloned in lentiCRISPRv2 plasmid using BsmBI restriction site. Sequence of the guide RNAs were as follows – PEX5 Forward 5’CACCGCACCATGGCAATGCGGGGAGC3’, PEX5 Reverse 5’AAACGCTCCCGCATTGCCATGGTGC3’, Scrambled Forward 5’CACCGCGGGACGTCGCGAAAATGTA3’, Scrambled Reverse 5’AAACTACATTTTCGCGACGTCCCGC3’. For Lentivirus production, cloned lentiCRISPRv2 plasmid was transfected in HEK293T cells together with helper plasmid pMD2G and psPAX2 with the help of PEI-MAX 40000. Post 72 hours of transfection, cell culture medium containing lentiviral particles were collected for consecutive three days and concentrated using PEG8000. For generation of knockout 50% confluent cells mixed with lentiviral supernatant into culture medium in 1:1 ratio along with polybrene (8 µg/ml). After 24hrs of transduction, the media (DMEM) was changed containing with 10% FBS/antibiotic. Then, cells were incubated in media with 2ug/ml puromycin for selection. For PEROXO-Tag IP studies, a lentiviral plasmid pLJC5-3XMyc-EGFP-PEX26) was directly purchased from Addgene (Plasmid, #139059). Lentivirus production and transduction of HeLa cells were done similarly as mentioned above.

### RNA extraction and quantitative PCR

The total cell RNA from samples was isolated using TRIzol (Ambion). Reverse transcription was carried out using Taqman reverse transcription kit (Applied Biosystems). Peroxisome and trafficking pathway gene primers were designed by NCBI and purchased from Sigma. The housekeeping gene GAPDH was used as normalizing control to calculate the fold change.

### Western blotting

Cells were lysed in lysis buffer (RIPA buffer) containing protease inhibitor cocktail. Then the cell lysates were separated by using SDS-PAGE and was transferred onto PVDF membrane (BioRad). The membrane was blocked using 5% non-fat dried milk. After washing with TBST (Tris-base, Nacl, pH -7.6 with Tween-20), the membrane was then incubation with primary antibody overnight at 4°C and HRP conjugated secondary antibody for 2 hours at room temperature. The immunodetection was achieved using chemiluminescence. The bands were quantified using ImageJ software (1.53k).

### Gentamicin protection assay

Cells were infected with STM WT or STM *ΔsseI* or STM at MOI of 10 (for intracellular survival assay) and MOI 20 (for qRT-PCR and immunofluorescence). After infection, plate containing infected cells was centrifuged at 600 rpm for 10 minutes to facilitate the proper adhesion. The plate was then incubated for 25 minutes at 37°C humidified chamber and 5% CO2. Then the media was removed from each well and the cells were washed with 1X PBS. Fresh media containing 100 µg/mL gentamicin was then added and again incubated for 60 minutes at 37°C humidified chamber and 5% CO2. The media was then removed, and cells were washed with 1X PBS twice and fresh media containing 25µg/mL gentamicin was added. The plate was incubated for at 37°C humidified chamber and 5% CO2 till the appropriate time. For the intracellular survival assay the cells were incubated till 2 hours and 16 hours post infection.

### Intracellular survival assay and invasion assay

Following gentamycin protection assay, the cells were lysed using 0.1% Triton X followed by addition of more 1X PBS at the appropriate time points (2 hours and 16 hours post infection). The collected samples from each well were plated at the required dilutions on LB agar plates and kept at 37°C for 12-14 hours. Post incubation the colony forming units (CFU) were enumerated for each plate. The fold proliferation and percentage invasion were determined as follows:

Fold Proliferation = CFU at 16 hours / CFU at 2 hours

Percentage Invasion = [CFU at 2 hours/ CFU of the pre-inoculum] ×100

### In-vivo animal experiment

5-6 weeks old C57BL/6 mice were infected orally by gavaging 10^7^ CFU of STM WT, STM *ΔsseI* (n=6). 7 days post-infection, the mice were sacrificed, and intestine, mesenteric lymph node (MLN), spleen and liver were isolated aseptically. The tissue samples were crushed using 1mm glass beads in a bead-beater, and the supernatant was plated onto SS agar and incubated at 37°C. CFU were enumerated after 16 hours, and organ load was determined by normalising the CFU to weight of the tissue sample.

### Bacterial gene knock-out generation

The gene knock-out in bacteria was done using the One-step chromosomal gene inactivation protocol by Datsenko and Wanner (2000). Briefly, primers were designed for amplification of Kanamycin resistance gene cassette from PKD4 plasmid. The 5’terminus of the primers had sequence homologous to the flanking region of the gene to be knocked out (here *sseI*). After amplifying the Kanamycin resistance gene cassette, the amplified products were purified using chloroform-ethanol precipitation. The purified product was then electroporated into the STM WT strain (expressing PKD-46 plasmid to provide λ-Red recombinase system) by a single pulse of 2.25 kV. Immediately fresh recovery media was added, and it was then incubated at 37°C for 60 minutes in an orbital-shaker. After incubation, the cultures were centrifuged at 8000rpm for 6 minutes. Then the pellet was dissolved in 100μL of media and plated at the required dilution on LB agar plates with Kanamycin. The plates were incubated at 37°C for 12-14 hours. The colonies were selected and were confirmed for knock-out using PCR which was then run on 1% agarose gel to compare the length of the products in the mutant from STM WT bacteria.

### Pip2 ASSAY

HeLa cells infected with STM, STM *ΔsseI*, siRNAs and inhibitors for different time point were checked for PIP2 level by using PIP2 Mass ELISA kit (Echelon Biosciences Inc., USA K-4500).

### Cholesterol assay

HeLa cells (PEROXO-tagged) were infected with STM, STM *ΔsseI* and siRNAs for different time point. And then, cholesterol level was measured from isolated peroxisome from those cells with the Amplex Red Cholesterol Assay kit (A12216). For the fluorometric quantification, isolated peroxisome and cholesterol standards were incubated in 96-well plates according to the instructions of the manufacturer kit (A12216).

### SseI construct and mutant generation

SseI gene was amplified from *Salmonella typhimurium* (STM) strain LT-2 strain using high fidelity DNA polymerase enzyme from TaKaRa and cloned in with N-terminal flag and HA tag in mammalian expression vector pcDNA3 flag HA Akt1 plasmid obtained from addgene(1477). Various SseI mutants (GKM mutant K321A) included in our study were generated using Agilent Quikchange site directed mutagenesis kit ().

### ARF1 clone and mutant generation

Gene encoding ARF-1 were amplified from cDNA (cDNA synthesized from RNA isolated from Hela cells) using high fidelity DNA polymerase enzyme from TaKaRa and cloned with N-terminal flag and HA tag in mammalian expression vector pcDNA3 flag HA Akt1 plasmid obtained from addgene(1477). Analysis of ARF-1 mediated trafficking was done by ARF-1 GTP bound mutant ARF1Q71L. Arf-1 cloned in above vector was subjected to site-directed mutagenesis (ARF1Q71L) using Agilent Quikchange kit.

### Peroxisome isolation by PEROXO-Tag IP

Cells (∼4 million HeLa cells) were transfected with three <u>Myc</u> epitopes (3XMyc- EGFP-PEX26). Then cells were rinsed twice with pre-chilled PBS and then scraped in 1 ml of KPBS and pelleted at 1000 x g for 2 min at 4°C. Cells were then resuspended in 1000 µl of KPBS and gently homogenized with 25 strokes for 10s of a 2 ml homogenizer. It was then centrifuged at 1000 x g for 2 min at 4°C to pellet nuclei and cells while cellular organelles including peroxisomes remained in the supernatant which was incubated with anti-Myc antibody at 4°C overnight on rocker. Next day, it was incubated with protein A beads for 4h at 4°C and was centrifuged at 1000xg for 2 min. The bound peroxisomes were resuspended in 1X SDS lysis buffer to extract the proteins.

### Catalase activity

HeLa cells infected with STM for different time points (3hr, 6hr) were checked for catalase activities by using the BioAssay Systems EnzyChrom catalase assay kit (Universal Biologicals Ltd., Cambridge, UK), according to the manufacturer’s instructions.

### SseI protein purification and SseI-ARF1 in vitro and in vivo binding

SseI Protein purification: SseI (322 residue) was cloned from genomic DNA of STM strain LT2 into pet23a vector. The construct was transformed into the *Escherichia coli* BL21 (DE3) strain and grown in LB medium at 200 rpm to stationary phase followed by induction with 1 mM IPTG. Culture was then grown overnight at 16°C at

200 rpm. The cells were harvested by centrifugation, resuspended in 20 mM Tris pH 8.0, 150 mM NaCl,0.3% triton X 100,5% glycerol and 1 mM PMSF (added during lysis) and lysed using sonicator. After centrifugation, the lysates were loaded onto a gravity-flow column containing Ni–NTA resin (Qiagen) equilibrated in lysis buffer (without PMSF). This was followed by a wash (wash buffer containing 20mM tris buffer PH-8.0, 300mM NACl,30 mM imidazole and 1mM PMSF) and protein elution (Elution buffer containing 20mM tris buffer PH-8.0, 50mM NACl,300mM imidazole and 1mM PMSF) was done. Following overnight dialysis into 50 mM Tris pH 8.0, 200 mM NaCl, 2 mM DTT. Elute fractions were separated using SDS-PAGE.

Overexpressed ARF1-HA cell lysates were incubated with purified His-sseI for 2hrs at 4°C. Then the samples were incubated with Ni-NTA beads overnight at 4°C. The beads were washed, and bound proteins were eluted by boiling in SDS-sample buffer. All elutes were subjected to SDS-PAGE and coomassie staining or immunoblotting.

IP experiments:

Cells transfected with the plasmids were lysed with RIPA lysis buffer containing protease inhibitor cocktail. The cell lysates were incubated overnight at 4°C with the respective primary antibodies (control- IgG). The antibody-bound proteins were pulled down by protein A beads and subjected to immunoblotting analysis.

### Statistical analysis

Statistical analyses were performed in GraphPad Prism software v7. Significance was referred as Student T-test, ANNOVA, *, **, *** and **** for P-values < 0.05, < 0.01, < 0.001 and < 0.0001 respectively.

## Acknowledgements

The work is supported by financial grants from the Science and Engineering Board (SERB), Government of India (SRG/2019/000268), ICMR Ad-hoc(Ortho) 20202-NCD-1, CSIR-MLP 2105, CSIR HCP-0047 and start-up funding from Director CSIR-CDRI to AL and PDF/2021/002843 to VA. DR, JS, SP receives fellowship from CSIR, and SS receives fellowship from UGC, government of India. We would like to thank technical help from Mrs. Reema Roy, Dr. C.P. Pandey and Mr. Anil Verma for microscopy and Dr. AL Vishwakarma for Flow cytometry studies and Sophisticated Analytical Instrument Facility and Research Division, CSIR-Central Drug Research Institute. CSIR THUNDER (BSC0102) and MOES (GAP0118) Intravital facility is acknowledged. We thank Dr. Anil Gaikwad, Dr. Sachin Kumar and Dr. Dipak Dutta for helping us with reagents. This manuscript has a CSIR-CDRI communication number: 13/2023/AL.

## Author contribution

D.R., A.V.N., J.S., S.P., A.K., S.B., S.S., S.S. P.R., V.N., designed, performed and analysed the experiments. V.B., T.C., U.C.G, A.D., M.I.S., S.K.G., D.C., V.N., and A.L. designed and analysed data. D.C., V.N., and A.L. wrote the manuscript, obtained funding. A.L. steered the study.

---

**Supplementary Figure 1.**
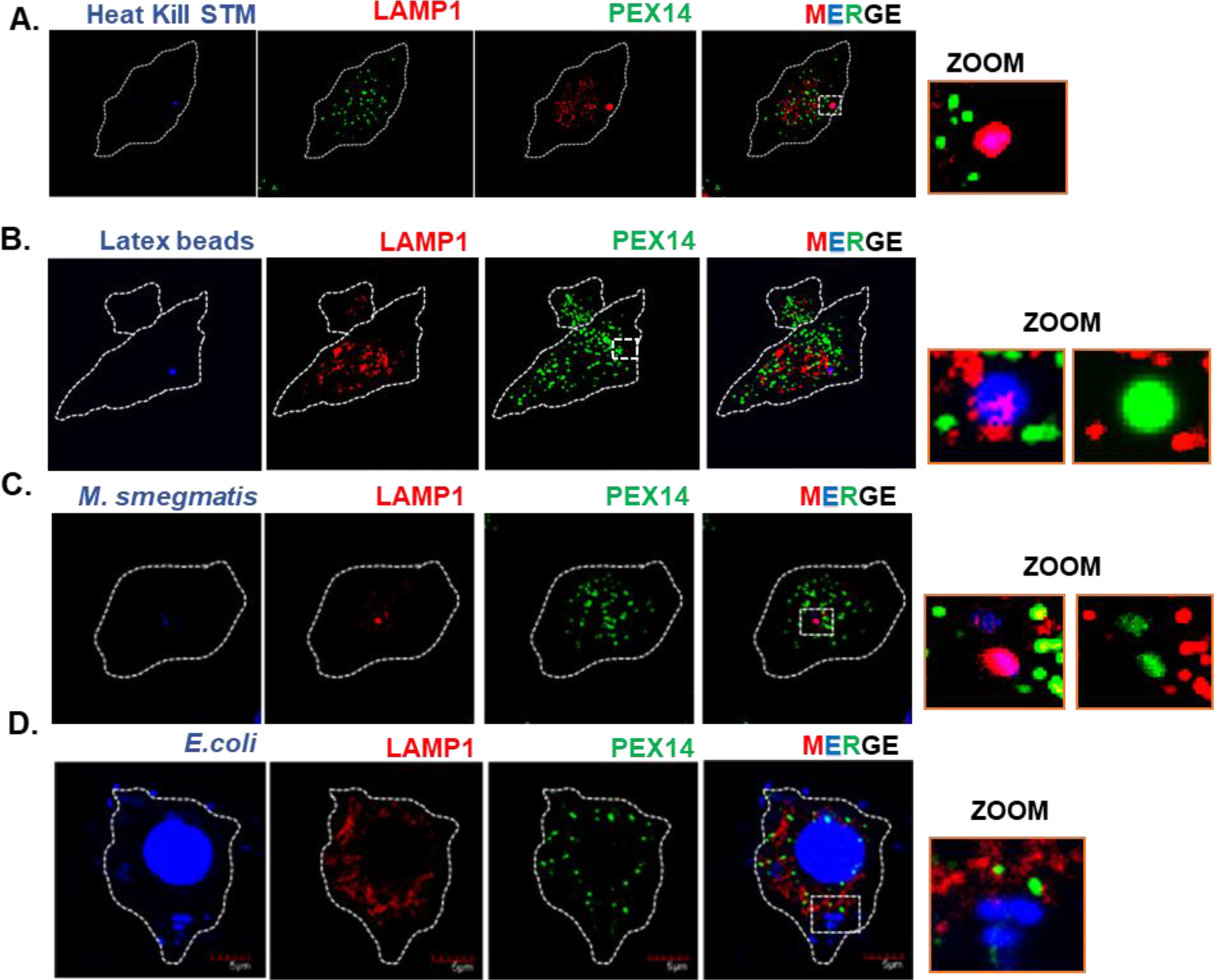
Peroxisomes do not interact with SCV after infection with heat killed STM and other bacteria. To understand peroxisomal role (dynamics and function) human monocyte derived macrophages (MDMs) were infected with A. heat killed STM, B. latex beads, C. *Mycobacterium smegmatis* and D. *E.coli* for 6 hours. After that cells were fixed, stained with Pex14 (green) and LAMP1 (red). heat-killed STM (blue), latex beads (blue), *Mycobacterium smegmatis* (blue)*, E.coli* (DAPI-blue). In all images, zoom part shows colocalization with peroxisome was absent. Scale bars 10 um.

**Supplementary figure 2.**
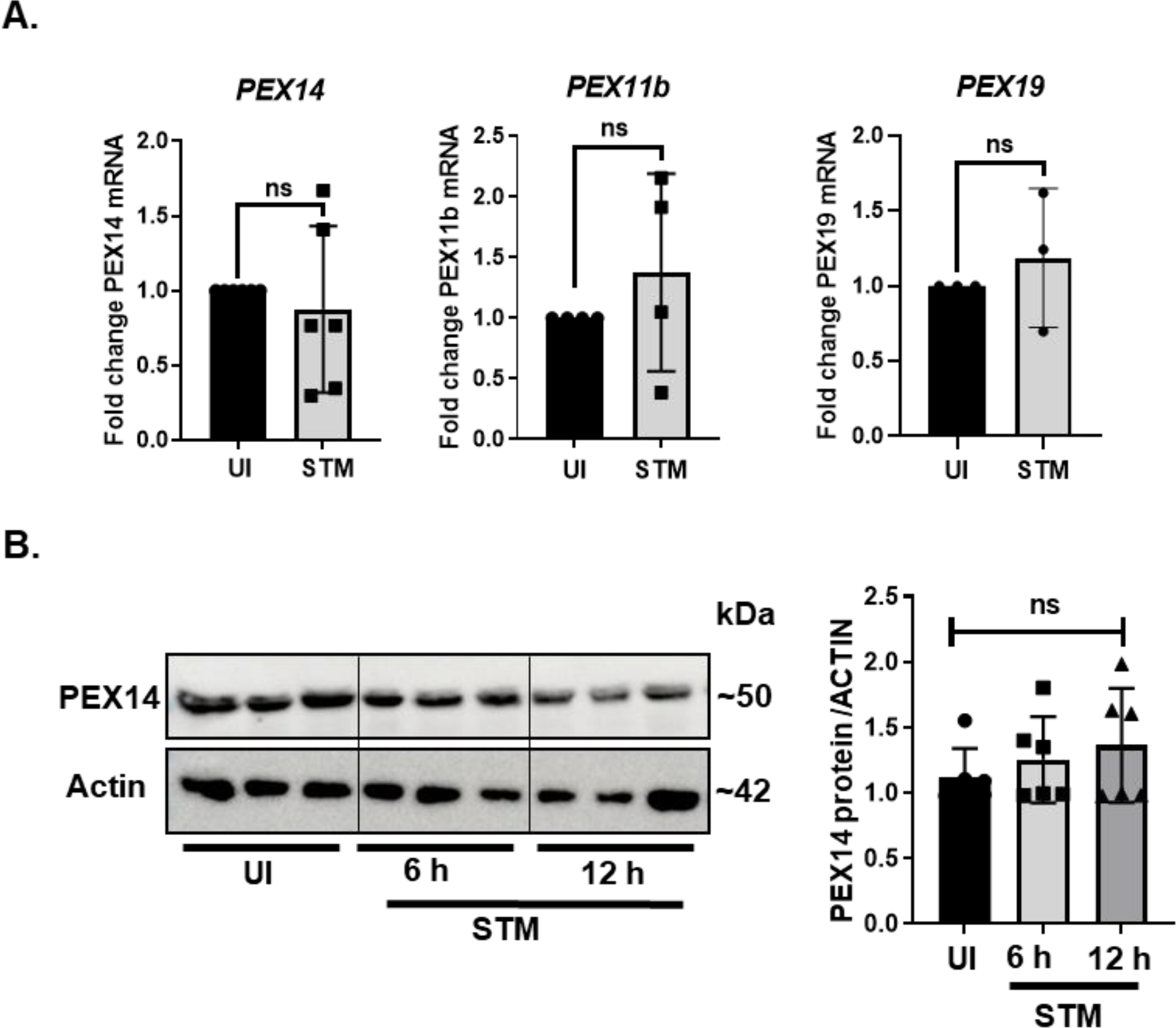
Peroxisome biogenesis, fission and fusion associated proteins are not altered after STM infection. A. Fold change in mRNA levels of indicated peroxisomal genes (pex14, pex11b and pex19) in THP1 cells post 6 hours of STM infection. B. Representative immunoblots of Pex14 and actin from whole cell lysates (HeLa) after STM infection for 3, 6 and 12 hours. Densitometry analysis denotes the abundance of Pex14 relative to Actin. Individual data points represent mean ±SD. ‘ns’ denotes non-significant, unpaired two-tailed Student T-test.

**Supplementary figure 3.**
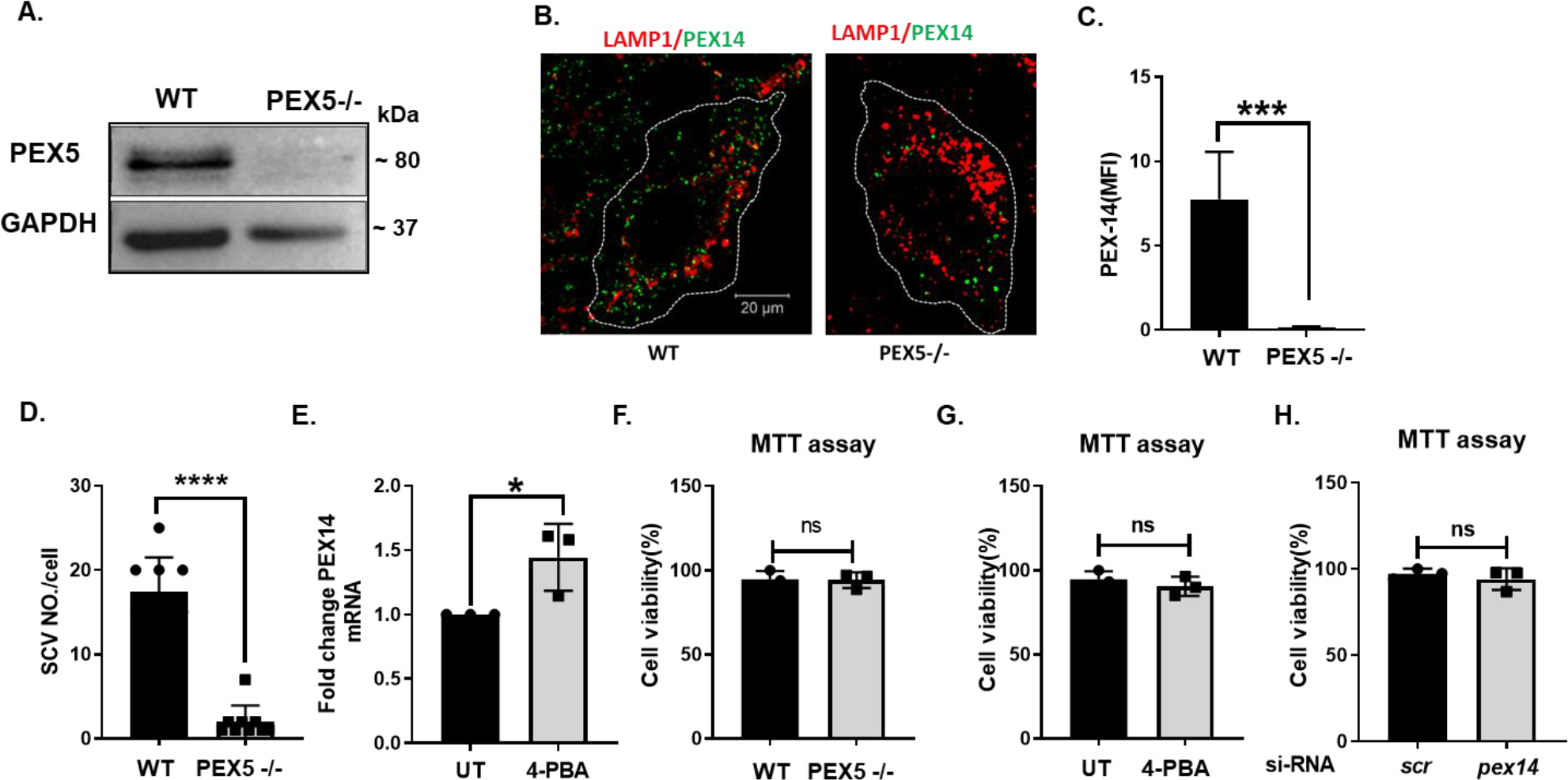
Validation of PEX5 KO HeLa cells. A. PEX5 KO HeLa cells were made by the CRISPR/Cas9 system. PEX5 Knock out efficiency was confirmed using western blot. B. Representative microscopy images of WT and PEX5 KO HeLa cells which were stained with LAMP1(red) and Pex14(green). The mean fluorescent intensity (MFI) quantification was presented.; scale bar 10 um. C. Graph represented the change in number of SCVs per cells after 6 hrs of infection with STM in wild type and PEX5 KO HeLa cells. D. Pex14 expression was quantified by RT-PCR. E. Cell viability was checked by MTT assay in PEX5 knockout cells F. Cell viability was checked by MTT assay the cells treated with 4-PBA. Experiment was repeated atleast two times. Individual data points represent mean ±SD. Result is representative of 3 independent experiments. ‘MOI’ denotes multiplicity of infection. ‘ns’ denotes non-significant ****p<0.0001, ***p<0.001, *p<0.05, unpaired two-tailed Student T-test.

**Supplementary Figure 4.**
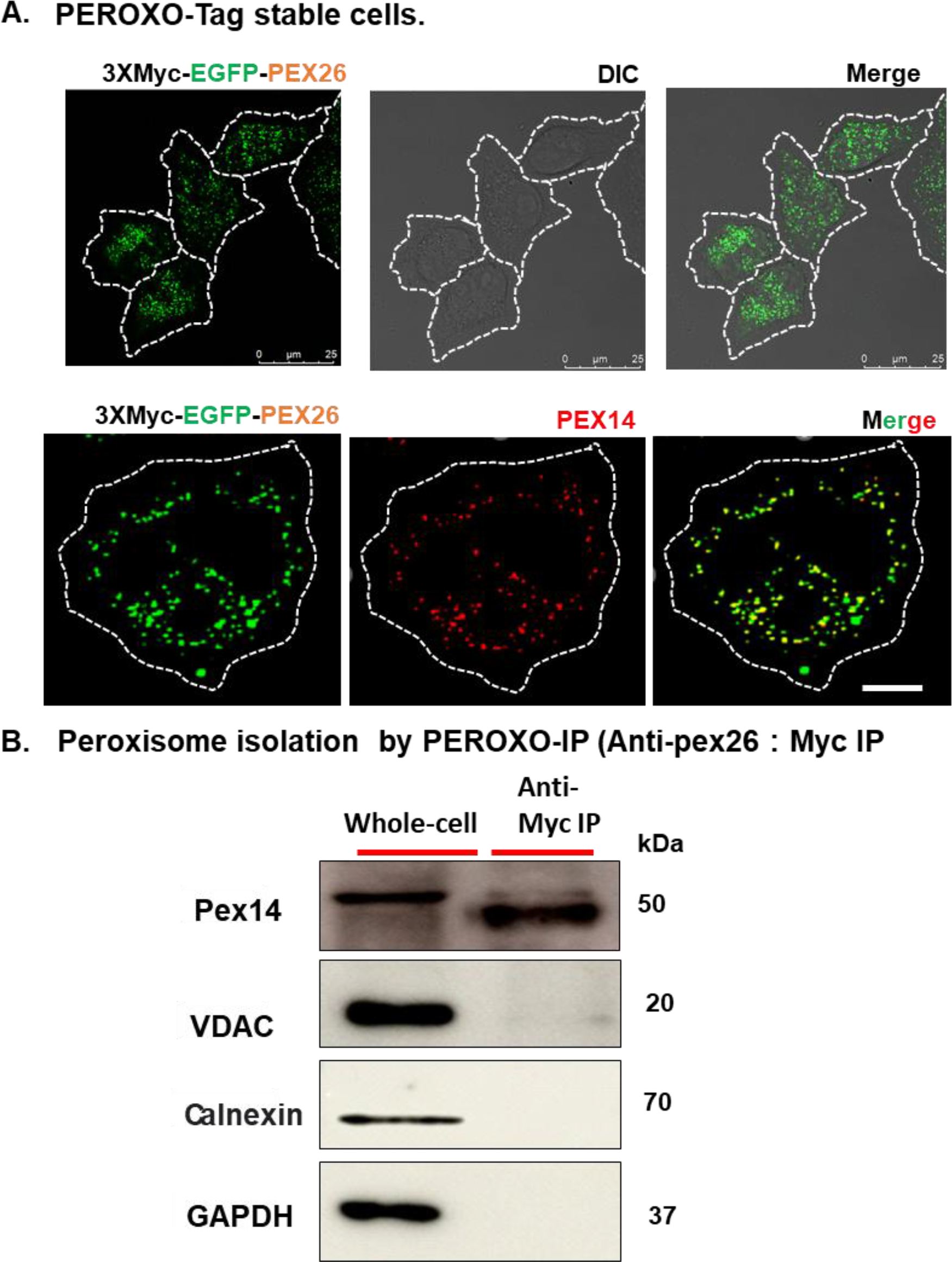
Generation of stable cell lines to study peroxisomal pathways. A. Representative microscopy images of PEROXO-tagged stable cell line where cells were transfected with lentivirus overexpressing 3X myc-EGFP-Pex26. Merge image represents co-localization of Pex14 with EGFP; scale bar 25 um. B. Purity of peroxisome isolated was confirmed by immunoblot stained with different organelle markers, Pex14 (for peroxisome), VDAC (for mitochondria), Calnexin (for ER) and GAPDH (for cytosol).

**Supplementary Figure 5.**
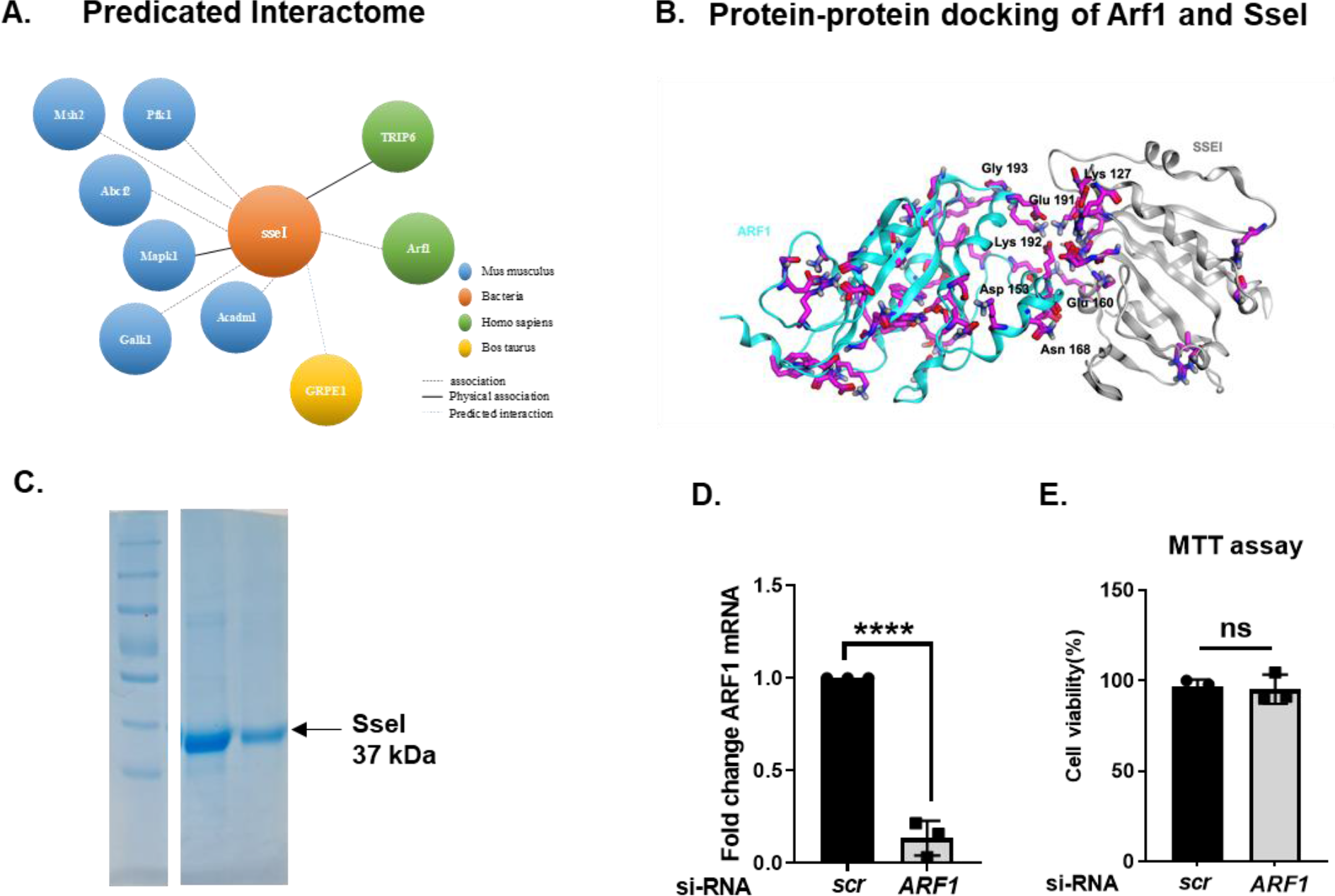
***In silico*** analysis to study the interaction between ***Salmonella*** effector protein, SseI and host GTPase ARF1. A. Predictive interactome of SseI from public database using Host Pathogen Interaction Database 3.0. B. Protein-protein docking of ARF1 with SseI subunit was performed. Docking score was calculated by using HDOCK SERVER. C. SseI: His purification (∼37 kDa) is shown. D. Knockdown efficiency of ARF1 in HeLa cells was validated using qPCR. E. Cell viability was checked by MTT assay in HeLa cells transfected with non-targeting siRNA (scr) and targeting ARF1. Individual data points represent mean ±SD. ‘ns’ denotes non-significant ****p<0.0001, unpaired two-tailed Student T- test.

**Supplementary figure 6:**
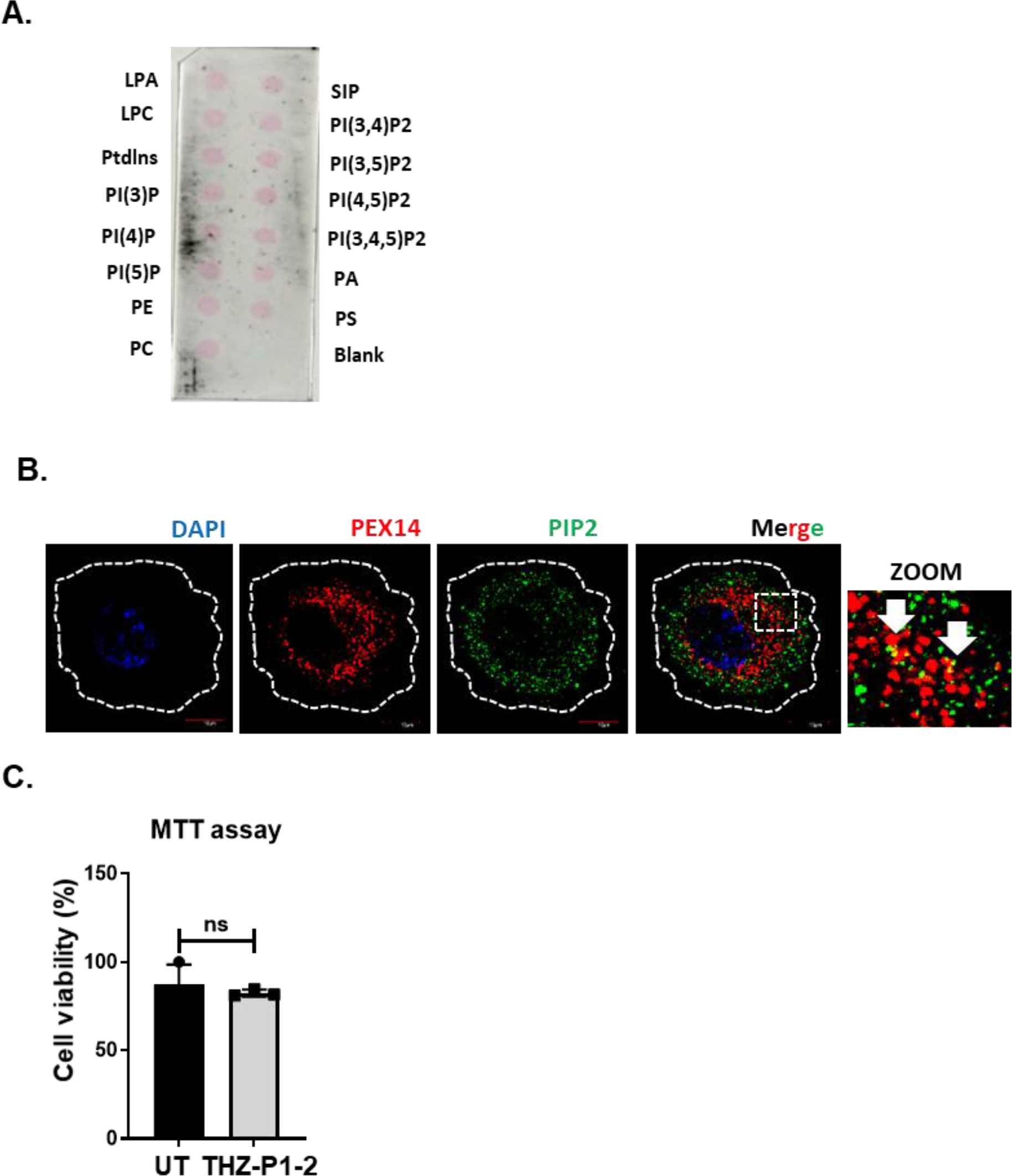
SseI do not interact with membrane lipids. A. Protein-lipid overlay was done to identify SseI and membrane lipid interaction. Purified SseI protein was incubated on lipid coated membrane. Scheme of the PIP-strip membrane is shown. Red line highlights the phospholipid species tested. B. Representative microscopy images of HeLa cells stained with Anti-PIP2 (green) and anti-PEX14 (red for peroxisome). Zoom part of image shows PIP2 (green), PEX14 (red for peroxisome) colocalization; (n=20 cells and similar results were found in independent experiment). Scale bar 10 um. C. Cell viability was checked by MTT assay in HeLa cells treated with THZ-P1-2. Individual data points represent mean±SD. Result is representative of 3 independent experiments. ‘MOI’ denotes multiplicity of infection. ‘ns’ denotes non-significant, unpaired two-tailed Student T-test.

**Supplementary figure 7:**
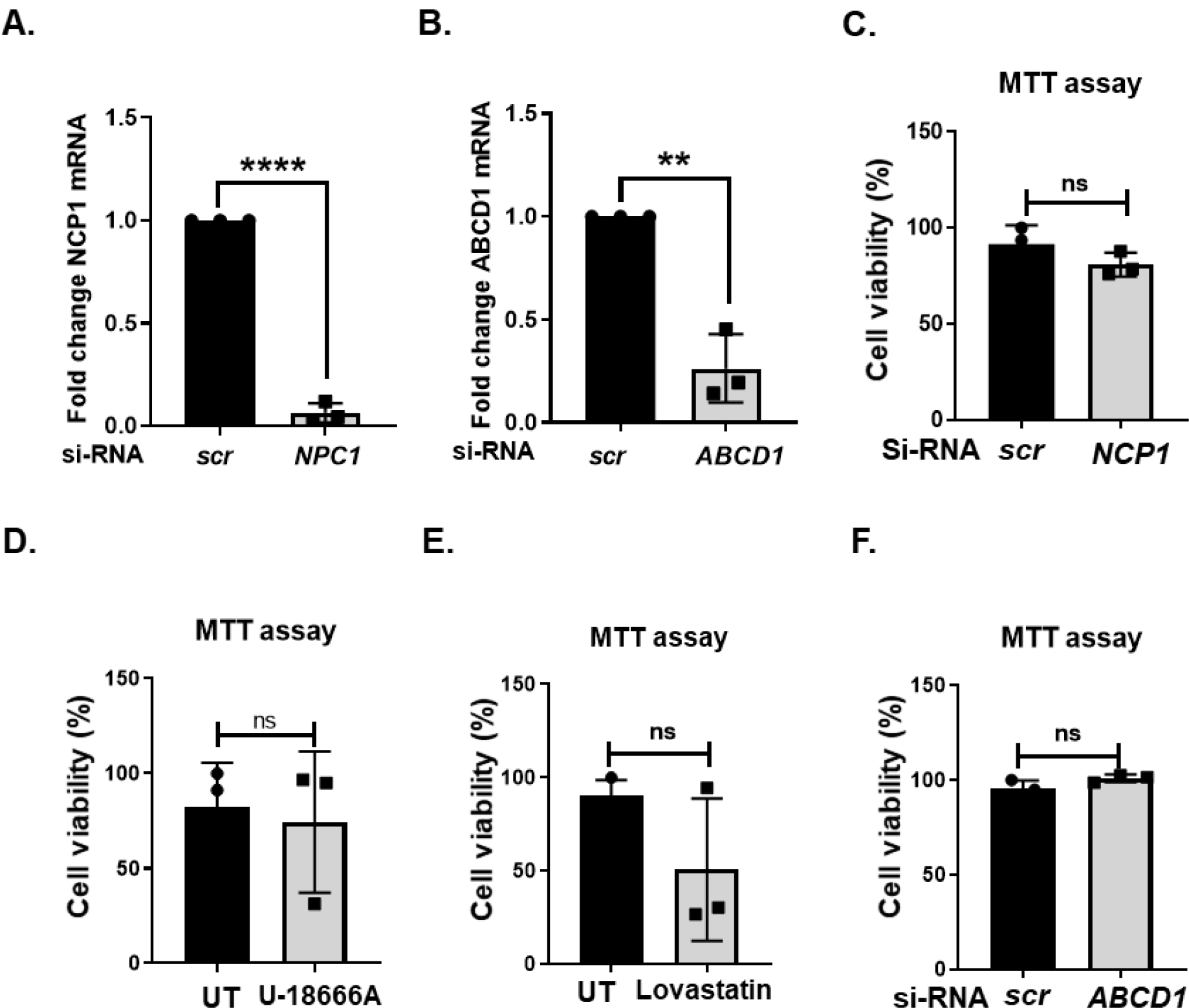
Knockdown efficiency and cell viability after knockdown. A-B. Graph representing the silencing efficiency of ABCD1 and NPC1 in HeLa cells. C-F. Cell viability was checked by MTT assay in HeLa cells transfected/treated with si-NPC1, U18666A, lovastatin and si-ABCD1. Individual data points represent mean± SD. Result is representative of 3 independent experiments. ‘MOI’ denotes multiplicity of infection. ‘ns’ denotes non-significant, **p<0.01 unpaired two-tailed Student T-test.

**Supplementary figure 8:**
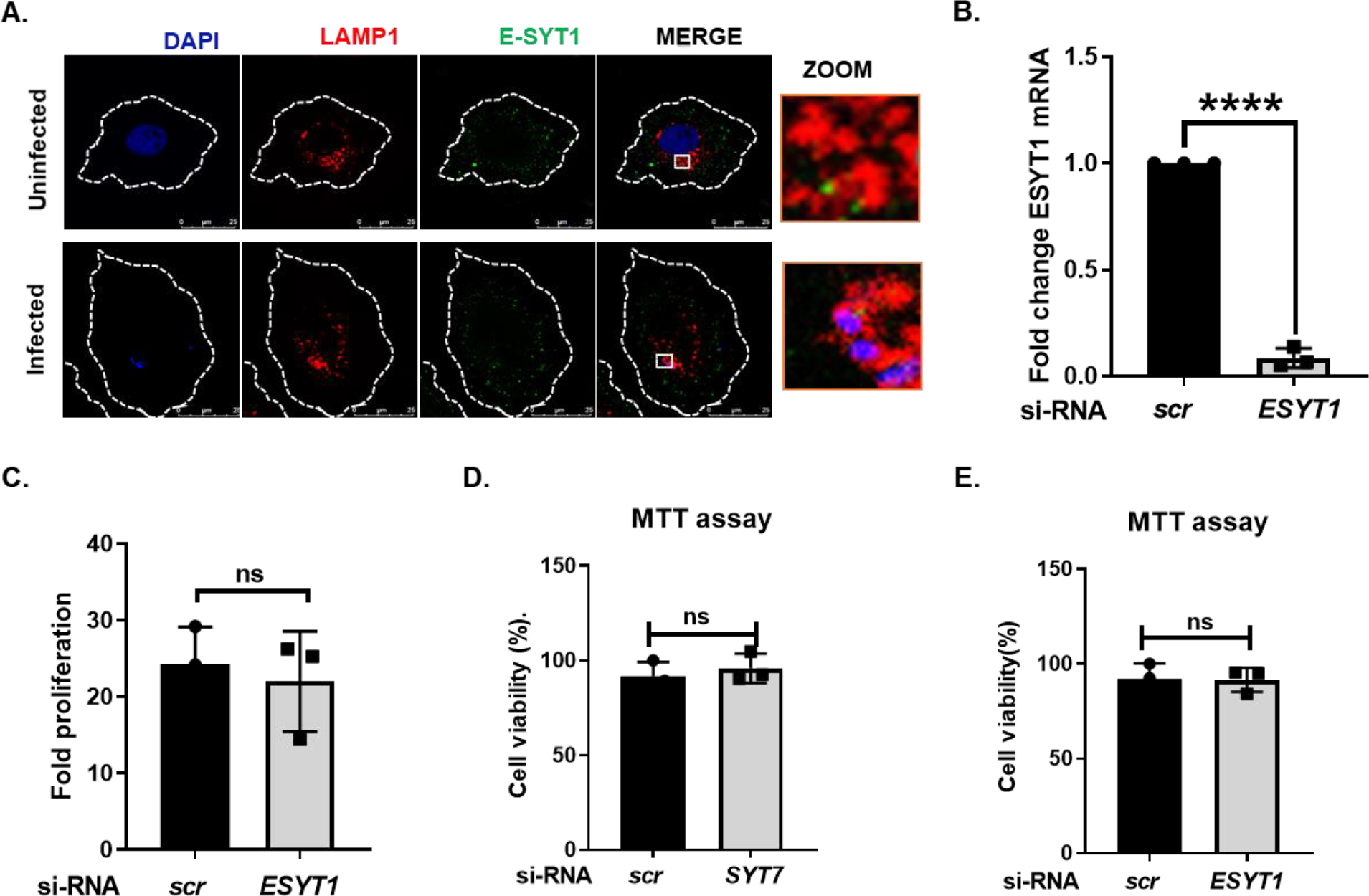
E-SYT1 is not recruited to SCV. A. Representative microscopy images showing staining of ESYT1 (green) and lysosome (LAMP1 red) in HeLa cells infected with STM (blue, DAPI) with MOI=50. Zoom part represented Lysosome (red), STM (blue) and ESYT1 (green) colocalization in the cells. Scale bar 25 uM. B. HeLa cells were transfected with non-targeting si RNA (scr) or si RNA against Esyt1 for 48 hr. Silencing efficiency of Esyt1 was measured by q-PCR.C. Changes in fold proliferation of WT STM after 2 and 16 hr post-infection is plotted in these cells. D. Cell viability was checked by MTT assay in HeLa cells transfected with non-targeting si RNA (scr) or si RNA against Esyt1 for 48 hr. Individual data points represent mean± SD. ‘ns’ denotes non-significant, ***p<0.001, unpaired two- tailed Student T-test.

**Supplementary figure 9:**
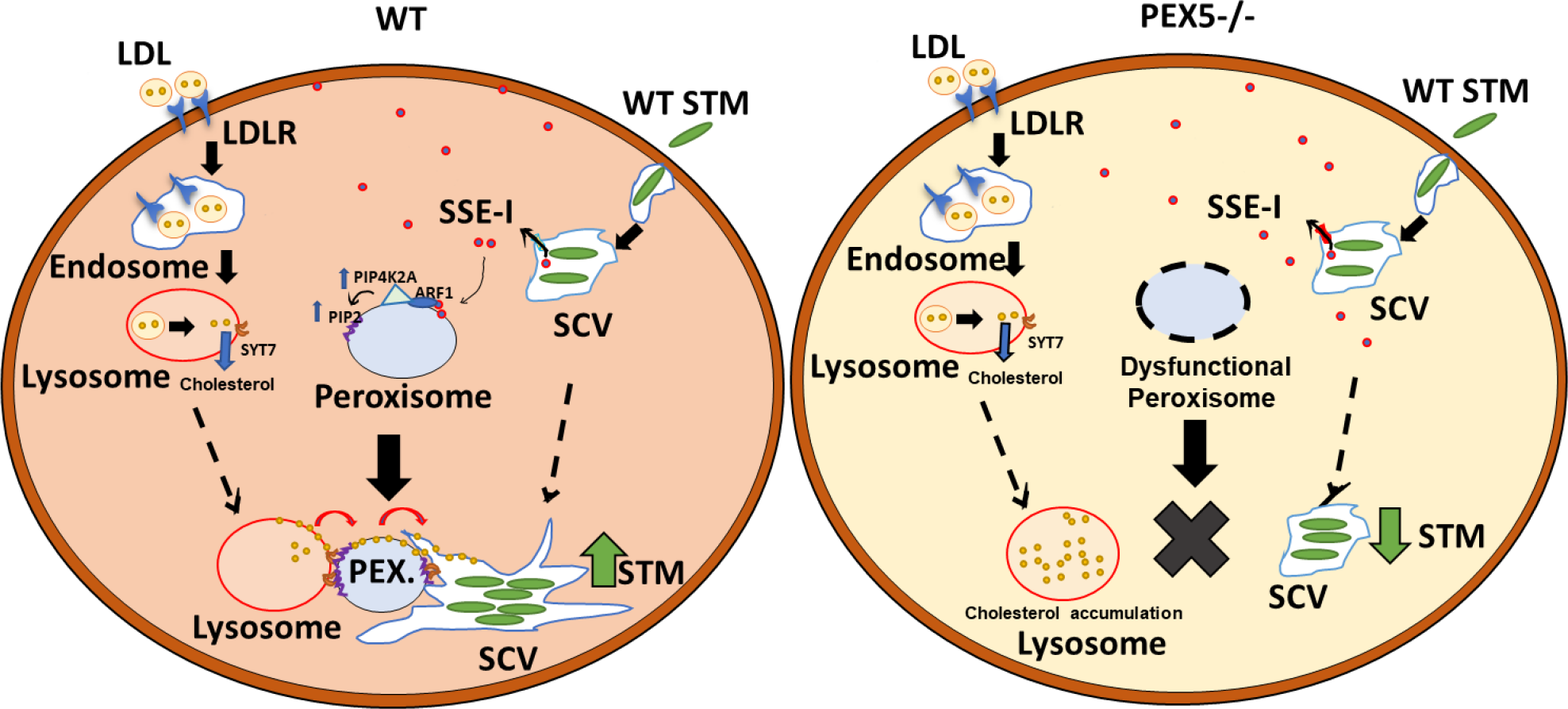
Schematic diagram. Mature SCVs secretes SPI-2 effector proteins in the host cell cytosol among which SseI contains host PTS1 signal. The effector protein, SseI interacts with GTPase ARF1 on host peroxisome. And recruits specific PIP4K2A which is essential for PIP2 synthesis on peroxisome. Simultaneously, cells receiving LDL from outside is further metabolized in lysosome and results in the release of free cholesterol. The lysosomal cholesterol is rerouted to the pathogenic bacteria using peroxisome as a bridge. This helps in SCV maturation and SIF formation resulting in the growth and proliferation of *Salmonella.* On the other hand, in the Pex5^-/-^ HeLa cells, dysfunctional peroxisome prevents its interaction with lysosome and SCV both. This results in the decreased proliferation of STM within the host cell.

